# PHF3 regulates neuronal gene expression through the new Pol II CTD reader domain SPOC

**DOI:** 10.1101/2020.02.11.943159

**Authors:** Lisa-Marie Appel, Vedran Franke, Melania Bruno, Irina Grishkovskaya, Aiste Kasiliauskaite, Ursula E. Schoeberl, Martin G. Puchinger, Sebastian Kostrhon, Etienne Beltzung, Karl Mechtler, Gen Lin, Anna Vlasova, Martin Leeb, Rushad Pavri, Alexander Stark, Altuna Akalin, Richard Stefl, Carrie Bernecky, Kristina Djinovic-Carugo, Dea Slade

## Abstract

The C-terminal domain (CTD) of the largest subunit of RNA polymerase II (Pol II) is a regulatory hub for transcription and RNA processing. Here, we identify PHD-finger protein 3 (PHF3) as a new CTD-binding factor that negatively regulates transcription and mRNA stability. The PHF3 SPOC domain preferentially binds to CTD repeats phosphorylated on Serine-2 and PHF3 tracks with Pol II across the length of genes. PHF3 competes with TFIIS for Pol II binding through its TFIIS-like domain (TLD), thus inhibiting TFIIS-mediated rescue of backtracked Pol II. PHF3 knock-out or PHF3 SPOC deletion in human cells result in gene upregulation and a global increase in mRNA stability, with marked derepression of neuronal genes. Key neuronal genes are aberrantly expressed in Phf3 knock-out mouse embryonic stem cells, resulting in impaired neuronal differentiation. Our data suggest that PHF3 is a prominent effector of neuronal gene regulation at the interface of transcription elongation and mRNA decay.

## INTRODUCTION

Transcription is a highly regulated process of RNA Polymerase II (Pol II) recruitment, initiation, pausing, elongation and termination (Chen et al., 2018; Core and Adelman, 2019; Kwak and Lis, 2013). Transcription regulation is critical to establish and maintain cell identity, and transcription misregulation underlies cancer and other diseases (Lee and Young, 2013). Transcription elongation factors modulate Pol II pause release, backtracking, elongation rate or processivity, and couple transcription elongation with co-transcriptional RNA processing (Chen et al., 2015; Close et al., 2012; Diamant et al., 2012; Fitz et al., 2018; Gregersen et al., 2019; Hou et al., 2019; Jonkers and Lis, 2015; Rahl et al., 2010; Saldi et al., 2016; Saponaro et al., 2014; Sheridan et al., 2019; Sims et al., 2004; Yu et al., 2015).

Transcription regulators interact with structurally defined surfaces of the Pol II complex and with the unstructured C-terminal domain (CTD) of the largest Pol II subunit RPB1 (Bernecky et al., 2017; Harlen and Churchman, 2017; Vos et al., 2018). Heptarepeats (Y_1_S_2_P_3_T_4_S_5_P_6_S_7_) within the Pol II CTD undergo dynamic phosphorylation during transcription, orchestrating the timely recruitment of regulatory factors. The early stages of transcription are marked by Pol II phosphorylated on Serine-5 (pS5), whereas productive elongation is characterized by the removal of pS5 and a concomitant increase in phosphorylated Serine-2 (pS2) (Harlen and Churchman, 2017). Transcription is tightly coupled with co-transcriptional RNA processing whereby Pol II CTD acts as a docking site for 5’ mRNA capping, splicing, 3’end processing, termination and mRNA export factors that recognize specific CTD phosphorylation patterns (Eick and Geyer, 2013; Herzel et al., 2017; Hsin and Manley, 2012; Saldi et al., 2016). 5’ mRNA capping enzymes employ the nucleotidyltransferase (NT) domain to directly bind pS5 within the Pol II CTD, whereas 3’end processing factors employ the CTD-interaction domain (CID) to directly bind the pS2 mark on the Pol II CTD (Doamekpor et al., 2014; Fabrega et al., 2003; Ghosh et al., 2011; Meinhart and Cramer, 2004).

The PHD-finger protein 3 (PHF3) belongs to a family of putative transcriptional regulators that includes the human Death-Inducer Obliterator (DIDO) and yeast Bypass of Ess-1 (Bye1) (Kinkelin et al., 2013; Wu et al., 2003). This family of proteins contains two motifs found in several transcription factors: a domain that is distantly related to the Pol II–associated domain of the elongation factor TFIIS, called the TFIIS-Like Domain (TLD), and a Plant Homeo Domain (PHD) (Kinkelin et al., 2013). It also contains a Spen Paralogue and Orthologue C-terminal (SPOC) domain, which has been associated with cancer, apoptosis and transcription (Sanchez-Pulido et al., 2004). Similar to TFIIS, Bye1 binds the jaw domain of RPB1 via its TLD *in vitro* and *in vivo* (Kinkelin et al., 2013; Pinskaya et al., 2014). In contrast to TFIIS, Bye1 TLD does not stimulate mRNA cleavage during transcriptional proofreading due to the absence of the TFIIS domain III (Cheung and Cramer, 2011; Kinkelin et al., 2013). Combined deletion of the PHD and SPOC domains abrogated Pol II binding by Bye1 *in vivo*, suggesting that the Bye1 TLD is necessary but not sufficient to interact with Pol II (Pinskaya et al., 2014). Although PHF3 does not contain any canonical CTD-binding domains, it was recently identified in a mass spectrometry screen for proteins binding to phosphorylated GST-CTD (Ebmeier et al., 2017). However, the physiological relevance of this interaction, and whether or how PHF3 regulates transcription, remain unknown.

Here, we discovered an unexpected interaction between PHF3 and the Pol II CTD via the SPOC domain. We show that PHF3 SPOC displays specificity towards CTD repeats phosphorylated on S2, establishing the SPOC domain as a new reader of the Pol II CTD. Moreover, we found that PHF3 exerts a dual regulatory function on both Pol II transcription and mRNA stability. While the SPOC domain enables docking of PHF3 onto Pol II, its TLD domain competes with TFIIS for Pol II binding. PHF3 thereby impairs TFIIS-mediated rescue of backtracked Pol II and negatively regulates transcription. Neuronal genes were aberrantly derepressed in PHF3 KO HEK293T cells, and Phf3 KO mESCs show impaired neuronal differentiation. Overall, our data suggest that PHF3, as a novel regulator of Pol II transcription and mRNA stability via the CTD, controls a neuronal gene expression program.

## RESULTS

### PHF3 interacts with RNA polymerase II through the SPOC domain

To explore the function of PHF3 in transcription, we expressed FLAG-PHF3 in HEK293T cells and identified interacting proteins by co-immunoprecipitation (co-IP) followed by mass spectrometry (Figure 1A,B). RPB1 ranked highest among 40 high-confidence PHF3 interactors, including other regulators of Pol II transcription elongation (SPT5, SPT6, PAF1C, FACT), as well as RNA processing factors (Figure 1A,B). We confirmed these findings by co-IP analyses of endogenous PHF3 tagged with GFP in HEK293T cells (Figure 1C,D and Figure S1A). Further, we found that endogenously expressed PHF3-GFP interacted with Pol II phosphorylated on Serines 2, 5 and/or 7 within the heptarepeats (Figure 1D,E).

**Figure 1.**
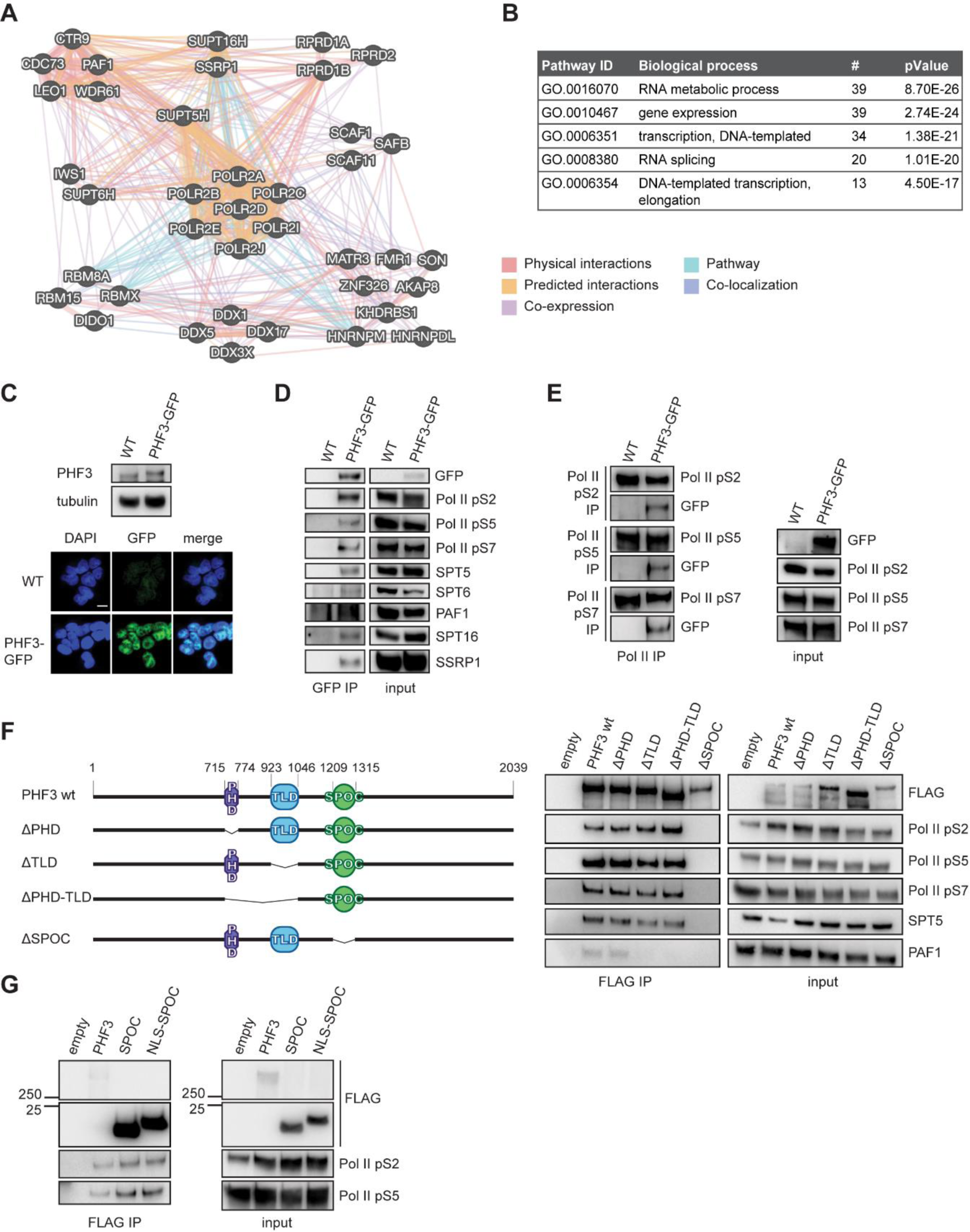
PHF3 interacts with RNA polymerase II via the SPOC domain. (A) GeneMANIA (Warde-Farley et al., 2010) interaction map of the PHF3 interactome. (B) Gene ontology biological processes of the PHF3 interactome revealed by mass spectrometry. (C) Expression levels and nuclear localization of PHF3-GFP in the endogenously tagged HEK293T cell line revealed by Western blotting with anti-PHF3 and fluorescence microscopy. Scale bar=10 μm. (D) Endogenously GFP-tagged PHF3 was immunoprecipitated using anti-GFP. Pol II phosphoisoforms, as well as transcription regulators SPT5, SPT6, PAF1 and FACT complex (SPT16 and SSRP1) were detected in the eluates. (E) Endogenous Pol II phosphoisoforms were immunoprecipitated from PHF3-GFP cells and PHF3-GFP was detected in the eluates. (F) Anti-FLAG immunoprecipitation of FLAG-PHF3 deletion mutants. Pol II does not co-immunoprecipitate in the absence of the PHF3 SPOC domain. (G) Anti-FLAG immunoprecipitation of the FLAG-SPOC domain shows interaction with Pol II.

To determine which domains within PHF3 are required for these interactions, we performed a co-IP analysis of different FLAG-PHF3 deletion mutants expressed in HEK293T cells (Figure 1F and Figure S1B,C). The truncation mutants localized to the nucleus and bound chromatin, similar to full-length PHF3 (Figure S1D,E). However, neither the TLD nor PHD domains were required for the interaction between Pol II and FLAG-PHF3 (Figure 1F and Figure S1C), which was unexpected given that the TLD is required for the Bye1-Pol II interaction in yeast (Kinkelin et al., 2013; Pinskaya et al., 2014). In contrast, removal of the SPOC domain from PHF3 completely abrogated the interaction with Pol II, the elongation factor SPT5 and the Pol II-associated factor PAF1 (Figure 1F). Moreover, the isolated PHF3 SPOC domain (aa 1199-1356) was sufficient to bind phosphorylated Pol II (Figure 1G).

### PHF3 SPOC preferentially binds RNA Pol II CTD phosphorylated on Serine-2

SPOC domains display low sequence conservation, but several amino acids within a positively charged patch on the SPOC surface are highly conserved, including an Arg residue and a Tyr residue (R1248 and Y1291 for PHF3 SPOC, Figure 2A). These residues are required for the SPOC-containing protein SHARP to interact with the co-repressor complex SMRT/NCoR (Ariyoshi and Schwabe, 2003). Serine phosphorylation of the LSD motif within SMRT or NCoR increases their binding affinity for the conserved Arg within the SHARP SPOC domain, suggesting that the SPOC domain is a phospho-serine binding module (Mikami et al., 2014; Oswald et al., 2016).

**Figure 2.**
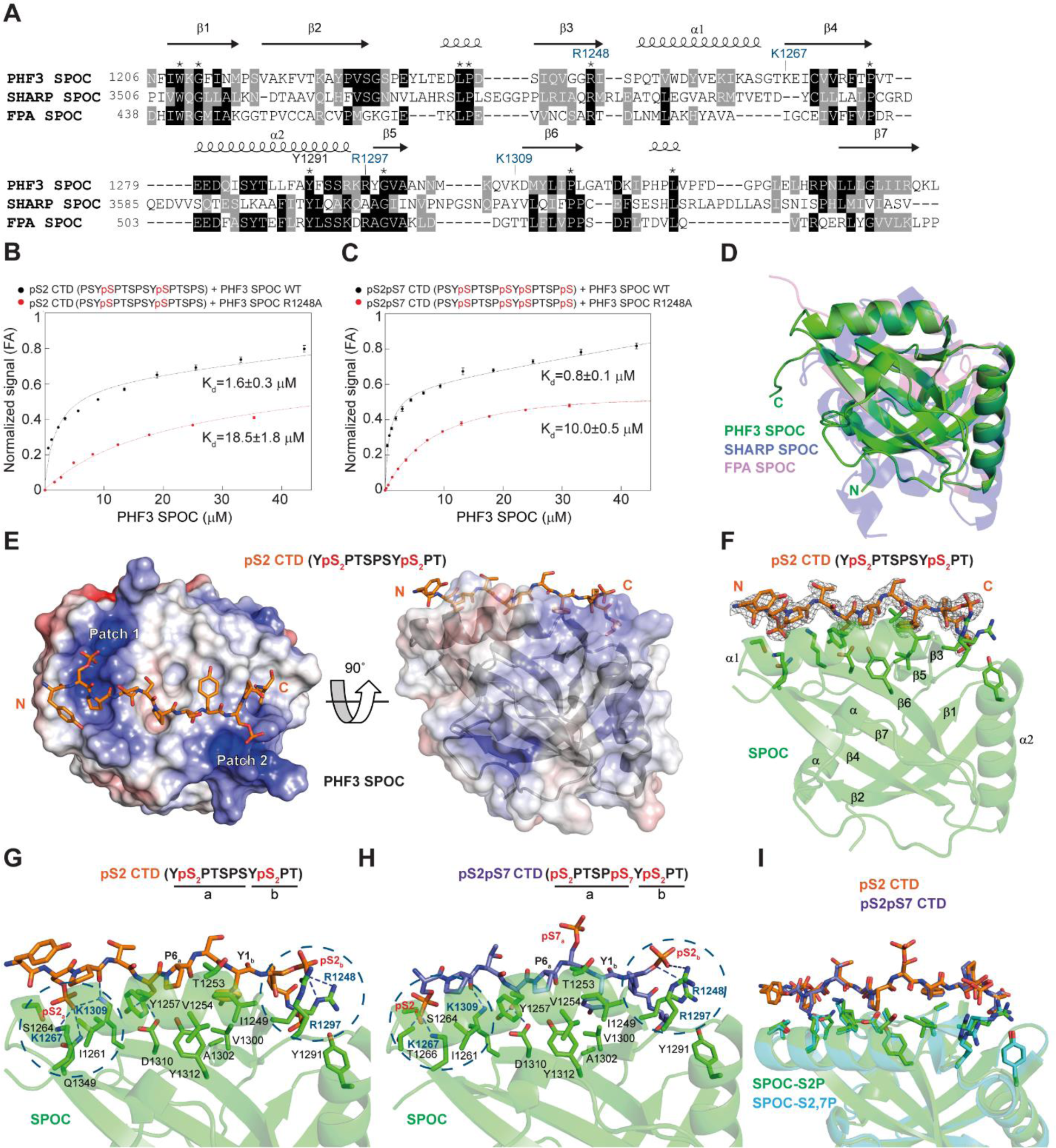
PHF3 SPOC binds pS2 CTD peptides *in vitro*. (A) Structure-based alignment of SPOC domains from PHF3 (6Q2V), SHARP (2RT5) and FPA (5KXF). Conserved residues are marked with an asterisk. (B,C) Fluorescence anisotropy (FA) measurement of the binding of (B) pS2 and (C) pS2pS7 CTD peptides to PHF3 SPOC WT or R1248A mutant. Normalized fluorescence anisotropy is plotted as a function of protein concentration (n=3). The data were normalized for visualization purposes and the experimental isotherms were fitted to one site saturation with non-specific binding model. (D) Overlay of SPOC structures from PHF3 (6Q2V), SHARP (2RT5) and FPA (5KXF) showed an average RMSD of 2.75 Å over 149 aligned Cα atoms between PHF3 and SHARP SPOC, and average RMSD of 1.94 Å over 123 aligned Cα atoms between PHF3 and FPA SPOC. (E) pS2 CTD peptide binds two positively charged patches (Patch 1 and 2) on the surface of PHF3 SPOC. The color coded electrostatic surface potential of SPOC was drawn using the Adaptive Poisson-Boltzmann Solver package within PyMol. The electrostatic potential ranges from −5 (red) to +5 (blue) kT/e. The N- and C-termini of the peptide are indicated and always shown in the same orientation. (F) 2 *F_o_* − *F_c_* electron density map of pS2 peptide contoured at the 1.5σ level. CTD peptide sequences used for X-ray structures correspond to those used in binding assays. Only the residues of the CTD diheptapeptide that are visible in the structure are indicated on the top. (G,H) Hydrogen bonding interactions between (G) pS2 and (H) pS2pS7 CTD peptides and PHF3 SPOC. SPOC monomer binds two phosphorylated S2 groups on the CTD peptides. SPOC residue labels from two positively charged patches are colored blue and the patches are contoured with dashed circles. (I) Overlay of SPOC co-structures with pS2 and pS2pS7 CTD peptides shows a similar extended conformation.

Based on these observations, we hypothesized that the positively charged surface of PHF3 SPOC binds the phosphorylated heptarepeats of Pol II CTD. To test this hypothesis, we examined the binding of bacterially expressed PHF3 SPOC to various phosphoisoforms of a CTD diheptapeptide (YSPTSPS-YSPTSPS) *in vitro*. PHF3 SPOC did not bind the unphosphorylated CTD diheptapeptide (Figure S2A), but phosphorylation of S2 within both repeats was sufficient to confer strong binding (Figure 2B,C and Figure S2A-E).

To uncover the basis of the CTD heptapeptide-SPOC interaction, we determined the structure of PHF3 SPOC in the apo form (2.6 Å), and bound to pS2 (2.0 Å), pS2pS7 (1.75 Å) and pS2pS5 (2.85 Å) diheptapeptides (Figure 2D-I, Figure S2I-L). Superimposition of PHF3 SPOC with SPOC domains from SHARP (Mikami et al., 2014) and FPA (Zhang et al., 2016) revealed that PHF3 SPOC has a distorted β-barrel fold comparable to other structurally characterized SPOCs, with seven β-strands and four helices connected with various loop regions (Figure 2D). The pS2- and pS2pS7-diheptapeptides were bound between strands of the β-barrel and along the α1 helix, in an extended conformation with *trans* isomer configuration of prolines (Figure 2E-I). The two phospho groups at S2 of the diheptapeptides were electrostatically anchored by two positively charged patches on the SPOC surface, which flank the central hydrophobic patch (Figure 2E-I and Figure S2H-L). SPOC binding to phosphorylated serines in adjacent repeats imposed an extended conformation on the CTD diheptapeptide (Figure 2E-I).

The ε-amino groups K1267 and K1309 from the first patch on SPOC form a hydrogen bond with O1P from the N-terminal pS2_a_, whereas guanidinium nitrogens from R1248 and R1297 from the second patch form hydrogen bonds with O1P and O3P from the C-terminal pS2_b_ (Figure 2G,H). Consistent with these findings, the R1248A substitution within SPOC reduced the binding to diheptapeptides containing pS2 more than 10-fold *in vitro* and reduced the interaction between PHF3 and Pol II in cells (Figure 2B,C and Figure S2B-F). pS7 within the pS2pS7 diheptapeptide did not form contacts with SPOC, as S7 was projected away from the SPOC surface (Figure 2H,I). I1249 and V1300 from the hydrophobic patch of SPOC form contacts with Y1_b_, and T1253, Y1257 and Y1312 form a hydrophobic pocket for P6_a_ (Figure 2G,H). All of these residues within PHF3 SPOC are generally conserved across species, as well as with DIDO SPOC (Figure 2A and Figure S2G,H). Fluorescence correlation spectroscopy (FCS) showed a 1:1 stoichiometry of the SPOC:pS-diheptapeptide interaction (Figure S3). Taken together, our data suggest that the PHF3 SPOC domain is a previously unrecognized Pol II CTD-binding domain that preferentially recognizes the elongating form of Pol II phosphorylated on S2.

### PHF3 travels with Pol II across the length of genes

Given that PHF3 binds pS2 Pol II, we hypothesized that they might co-localize throughout the genome. To examine their potential co-occupancy, we performed ChIP-seq of PHF3-GFP, Pol II pS2, Pol II pS5 and Pol II pS7. ChIP-seq analysis showed that PHF3 tracked with Pol II across the length of genes, with increasing strength from TSS (transcription start site) to pA (polyadenylation sites) (Figure 3A, Figure S4A). Additionally, higher PHF3 occupancy coincided with higher Pol II pS2 occupancy (Figure 3B and Figure S4B) and with higher transcription levels according to single base-resolution, precision nuclear run-on and sequencing (PRO-seq) (Figure S4C). Overall, these data indicate that PHF3 travels with the Pol II transcription machinery.

**Figure 3.**
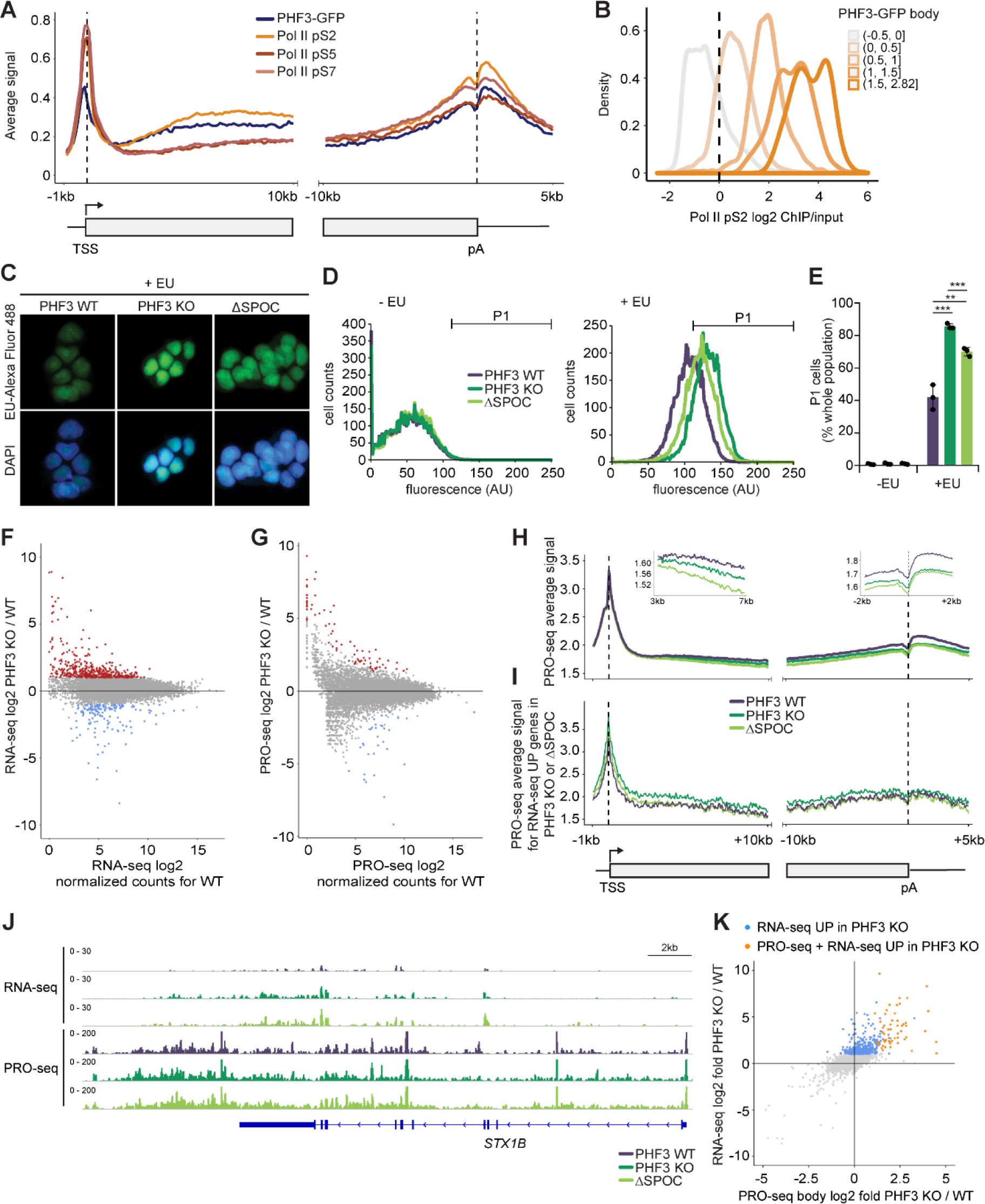
PHF3 travels with Pol II and negatively regulates transcription via the SPOC domain. (A) ChIP-seq analysis shows that PHF3 travels with Pol II across the length of genes. Relative enrichment of PHF3, Pol II pS2, pS5 and pS7 on TSS-gene body region (TSS viewpoint; left panel) and gene body-pA region (pA viewpoint; right panel) for genes that showed Pol II occupancy with the F-12 antibody (minimal gene body RPKM of 5 in F-12 ChIP-seq). (B) Pol II pS2 log2 ChIP/input in PHF3 WT colored by PHF3 gene body occupancy, with light color showing the lowest and dark color the highest PHF3 occupancy category. (C-E) PHF3 KO and ΔSPOC show increased incorporation of EU-Alexa Fluor^®^ 488 by (C) fluorescence microscopy and (D-E) FACS analysis. (D) Cell counts for each fluorescence intensity in the absence (-EU) or presence of EU (+EU). (E) Percentage of cells belonging to the gated P1 fluorescent population shown in (D) (n=3). (F) RNA-seq analysis shows upregulation of 620 genes (red dots, fold-change>2, p<0.05) and downregulation of 173 genes (blue dots, fold-change>2, p<0.05) in PHF3 KO compared to WT. Drosophila S2 cells were used for spike-in normalization. (G) PRO-seq analysis shows upregulation of 68 genes (red dots, fold-change>2, p<0.05) and downregulation of 39 genes (blue dots, fold-change>2, p<0.05) in PHF3 KO compared to WT. Drosophila S2 nuclei were used for spike-in normalization. (H) Composite analysis of PRO-seq distribution and signal strength in PHF3 WT, KO and ΔSPOC on TSS-gene body region and gene body-pA region for all genes. (I) Composite analysis of PRO-seq distribution and signal strength in PHF3 WT, KO and ΔSPOC on TSS-gene body region and gene body-pA region for RNA-seq upregulated genes (fold-change>2, p<0.05). (J) Genome browser snapshots showing RNA-seq and PRO-seq reads for *STX1B* as a typical gene with increased RNA-seq but no change in PRO-seq in PHF3 KO and ΔSPOC cells. (K) Relationship between RNA-seq and PRO-seq fold change for PHF3 KO vs WT. Genes that are upregulated in PHF3 KO in RNA-seq but not PRO-seq are indicated in blue. Genes that are upregulated in both RNA-seq and PRO-seq are indicated in orange.

### PHF3 negatively regulates gene expression

To elucidate the function of PHF3 and its SPOC domain in transcription, we used CRISPR/Cas9 to generate PHF3 knock-out HEK293T cells (PHF3 KO) and HEK293T cells lacking the SPOC domain (PHF3 ΔSPOC) (Figure S5). We initially assessed the effect of PHF3 loss or SPOC deletion by monitoring incorporation of EU (5-ethynyl uridine) during transcription (Figure 3C-E). EU was coupled to Alexa Fluor 488^®^ and visualized by confocal microscopy or quantified by FACS. PHF3 KO and PHF3 ΔSPOC cells showed significantly higher EU levels than WT cells (Figure 3C-E), suggesting increased transcript levels.

To investigate the changes in RNA levels genome-wide, we performed RNA-seq to measure mature transcripts, and PRO-seq to measure nascent transcripts (Figure 3F-K, Figure S6,7). We found that mature transcript levels were generally increased in both PHF3 KO and ΔSPOC cells compared to WT (Figure 3F and Figure S6A): 620 and 638 mature transcripts were elevated >2-fold (p-value<0.05) in PHF3 KO or ΔSPOC cells relative to WT, with 281 (∼45%) elevated in both conditions. In contrast, only 173 and 74 mature transcripts were downregulated >2-fold in PHF3 KO or ΔSPOC relative to WT, with 37 downregulated in both conditions.

Unlike mature transcripts, nascent transcripts did not show major changes in PHF3 KO and ΔSPOC cells (Figure 3G,H,J and Figure S6B). We observed increased nascent transcription of 68 and 78 genes >2-fold in PHF3 KO or ΔSPOC cells relative to WT, with 29 elevated in both conditions (Figure 3G and Figure S6B). Similarly, 39 and 70 genes showed decreased transcription >2-fold in PHF3 KO or ΔSPOC cells relative to WT, with 10 reduced in both conditions (Figure 3G and Figure S6B). Metagene profiles showed a small reduction in nascent transcript levels in the gene bodies and towards polyadenylation (pA) sites (Figure 3H).

About 10% of mature transcripts with elevated steady-state levels in PHF3 KO or ΔSPOC cells had a concomitant increase in nascent transcripts (Figure 3I,K). Overall, these data suggest that PHF3 acts as a repressor of gene expression, and that the binding of PHF3 to Pol II via the SPOC domain is necessary for its repressive function. PHF3 appears to exert its repressive effect by regulating Pol II transcription on a subset of genes (PRO-seq and RNA-seq upregulated, orange dots in Figure 3K) and by modulating mRNA stability on a global level (RNA-seq upregulated, blue dots in Figure 3K).

### PHF3 regulates transcription by competing with TFIIS

To understand how PHF3 may regulate Pol II transcription, we compared the Pol II interactome in PHF3 WT and KO cells, which revealed increased binding of transcription factors TFIIS and TFIIF to Pol II in PHF3 KO (Figure S8A). Pol II commonly arrests and backtracks within 300 bp of the TSS; TFIIS stimulates cleavage of the nascent RNA and allows transcription to continue (Cheung and Cramer, 2011; Izban and Luse, 1992; Nechaev et al., 2010; Sheridan et al., 2019). TFIIS and the TLD domain of the yeast PHF3 homolog Bye1 bind a similar region on Pol II (Kinkelin et al., 2013), suggesting that PHF3 competes with TFIIS for binding to Pol II. To test this hypothesis, we used sucrose gradient ultracentrifugation to analyze complexes formed between Pol II-EC (elongation complex) and PHF3 in the presence of TFIIS as a competitor, and vice versa (Figure 4A). We used an inactive TFIIS mutant (TFIIS^M^; D282A E283A) to prevent RNA cleavage and Pol II-EC disassembly. Although TFIIS could not displace PHF3 from Pol II-EC, PHF3 almost completely displaced TFIIS from Pol II-EC (Figure 4A). These data suggest that PHF3 competes with TFIIS for binding to Pol II.

**Figure 4.**
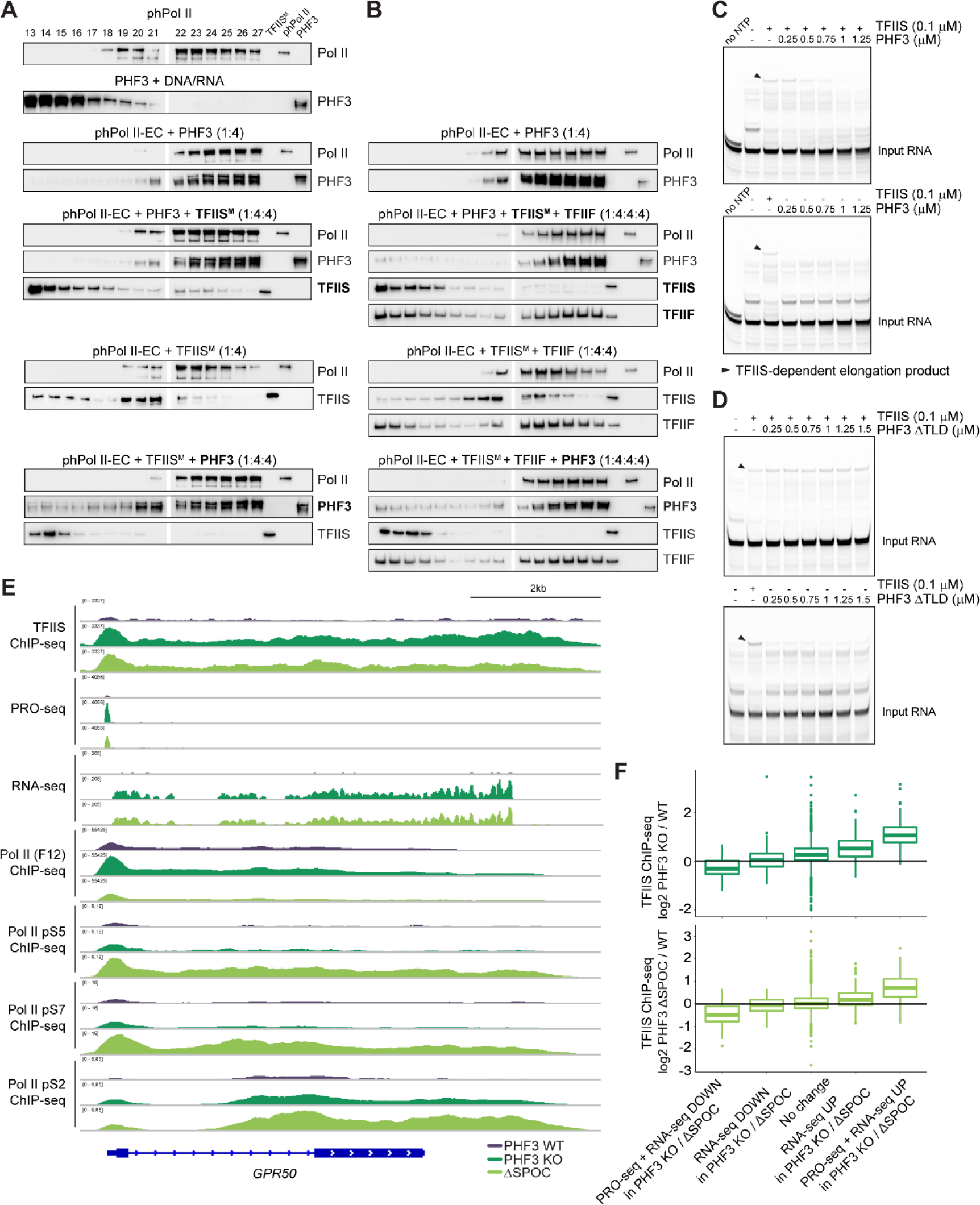
PHF3 negatively regulates transcription by competing with TFIIS. (A,B) Competition between PHF3 and TFIIS for complex formation with phosphorylated Pol II-EC. Pol II was phosphorylated with DYRK1A and Pol II-EC was formed using a DNA-RNA bubble scaffold. Complexes with Pol II-EC were preformed by adding 4-fold molar excess of PHF3 or (A) TFIIS^M^, (B) TFIIS^M^+TFIIF. A 4-fold molar excess of the competitor indicated in bold was added to the preformed complexes. Complexes were separated by sucrose gradient ultracentrifugation and fractions were analyzed by Western blotting. (C,D) *In vitro* assay monitoring Pol II elongation on an arrest sequence in the presence of TFIIS and increasing amounts of (C) PHF3 or (D) PHF3 ΔTLD (top gel) or in the presence of PHF3 alone (bottom gel). Pol II-EC was formed using an excess of a DNA-RNA bubble scaffold containing 5’-FAM-labeled RNA. The short elongation product seen in the ‘no NTP’ lane is due to residual ATP from the phosphorylation reaction. (E) Genome browser snapshots showing TFIIS ChIP-seq, PRO-seq, RNA-seq, and Pol II ChIP-seq (F-12, pS5, pS7, pS2) reads for GPR50. GAPDH as a housekeeping gene is shown in Figure S9. (F) TFIIS ChIP-seq log2 fold change PHF3 KO/WT (top) or PHF3 ΔSPOC/WT (bottom) for TSS. Genes were grouped according to changes in RNA-seq and PRO-seq: downregulation or upregulation in RNA-seq (fold change>2), downregulation or upregulation in RNA-seq and PRO-seq (fold change>2) or no change.

TFIIF, which cooperates with TFIIS to rescue arrested Pol II (Ishibashi et al., 2014; Schweikhard et al., 2014), stabilized the association between TFIIS and Pol II but did not prevent PHF3-mediated displacement of TFIIS (Figure 4B). TFIIF itself was not displaced by PHF3, nor did it outcompete PHF3 from the Pol II-EC (Figure 4B), indicating that PHF3 does not compete with TFIIF. Thus, even in the presence of TFIIF, PHF3 outcompetes TFIIS for binding to Pol II.

Furthermore, we tested whether PHF3 inhibits the function of wild-type TFIIS in a transcription assay using an arrest template (Figure 4C). Pol II-EC efficiently transcribed the arrest sequence in the presence of TFIIS, but elongation was diminished in the presence of PHF3 (Figure 4C). PHF3 lacking the TLD domain (PHF3 ΔTLD) did not interfere with TFIIS-dependent elongation, confirming that the PHF3 TLD domain is critical for competition with TFIIS (Figure 4D).

TFIIS ChIP-seq experiments showed increased TFIIS occupancy in PHF3 KO and ΔSPOC cells, in accordance with the competition between PHF3 and TFIIS (Figure 4E, Figure S8,9). Moreover, TFIIS occupancy was increased most prominently on genes that were both PRO-seq and RNA-seq upregulated in PHF3 KO or ΔSPOC, which also showed increased Pol II occupancy (Figure 4F and Figure S8).

Taken together, PHF3 negatively regulates transcription by competing with the transcription factor TFIIS and impairing rescue of backtracked Pol II. PHF3 competition with TFIIS explains the increase in both nascent and mature transcripts observed for a subset of genes in the absence of PHF3 or its SPOC domain.

### PHF3 regulates mRNA stability

The major phenotype of PHF3 loss or SPOC deletion was an increase in mature transcripts without major alteration in nascent transcription levels, suggesting that PHF3 regulates mRNA stability. To test this, we determined mRNA half-lives by performing SLAM-seq [Thiol (SH)-Linked Alkylation for the Metabolic sequencing of RNA] (Herzog et al., 2017) (Figure 5A). Cells were pulse-labelled for 12 h with s^4^U (0 h sample), followed by a chase with uridine for 6 and 12 h. The T-C conversion rate was used to determine mRNA half-lives for 755 genes, chosen because they had conversion rates >0 at the 0 h timepoint and showed a monotonic decrease in median conversion rates (Figure 5B, Figure S10).

**Figure 5.**
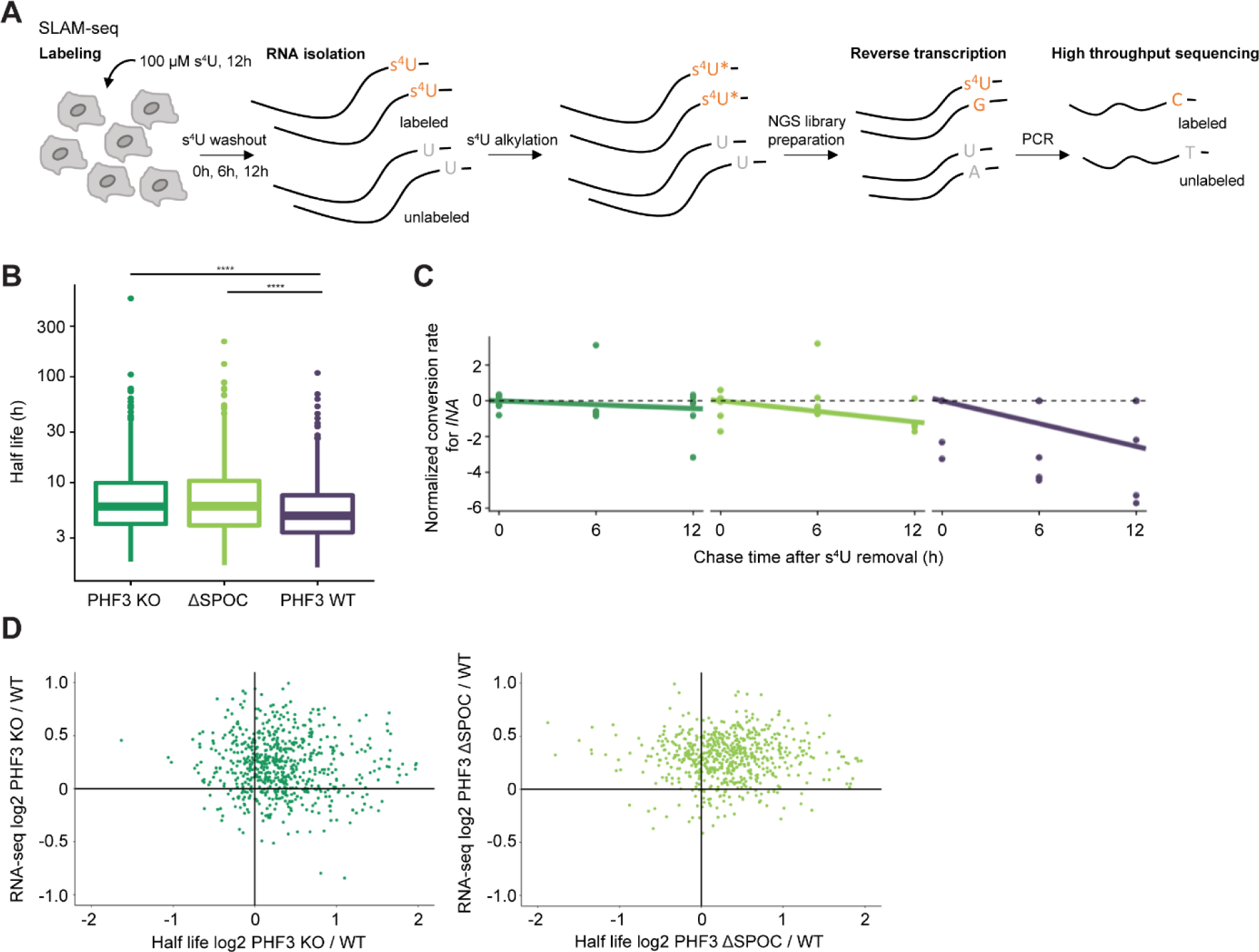
PHF3 negatively regulates mRNA stability via the SPOC domain. (A) Schematics for SLAM-seq. (B) Comparison of mRNA half-lives for 755 genes calculated from T-C conversion rates as determined by SLAM-seq in PHF3 WT, KO and ΔSPOC cells (n=6). The difference between the distributions is statistically significant based on the one-sided Wilcoxon test [*P*(KO – WT)=1.34x10^-11^, *P*(ΔSPOC – WT)=2.28x10^-11^]. (C) Conversion rates determined from targeted SLAM-seq analysis of *INA* mRNA labelled with s^4^U for 12 h followed by pulse chase for 6 h and 12 h. Robust linear models were fit on the linearized form of the exponential decay equation. Y-axis shows the log2 conversion rate, shifted by the median conversion rate at t = 0 h. (D) Relationship between RNA-seq fold change and half-life fold change for PHF3 KO vs WT (left) or PHF3 ΔSPOC vs WT (right).

We observed a significant overall increase in mRNA half-lives in PHF3 KO and ΔSPOC compared to WT (Figure 5B). Of the 755, 466 (62%) showed increased mRNA half-lives in both PHF3 KO and ΔSPOC, and an additional 206 (27%) showed an increased half-life in either KO or ΔSPOC. Because global SLAM-seq analysis failed to capture low-expressed genes, which exhibit the strongest upregulation in PHF3 KO and ΔSPOC (Figure 6D), we applied a targeted SLAM-seq approach to specifically analyze mRNA half-lives for *INA* as one of the most highly upregulated genes in PHF3 KO and ΔSPOC. *INA* showed a pronounced increase in mRNA half-life from 3.3 h in WT to 7.1 h in ΔSPOC and 19.5 h in PHF3 KO (Figure 5C). Comparison between RNA-seq and SLAM-seq data showed that increased mRNA stability correlated with mature transcript levels (Figure 5D).

**Figure 6.**
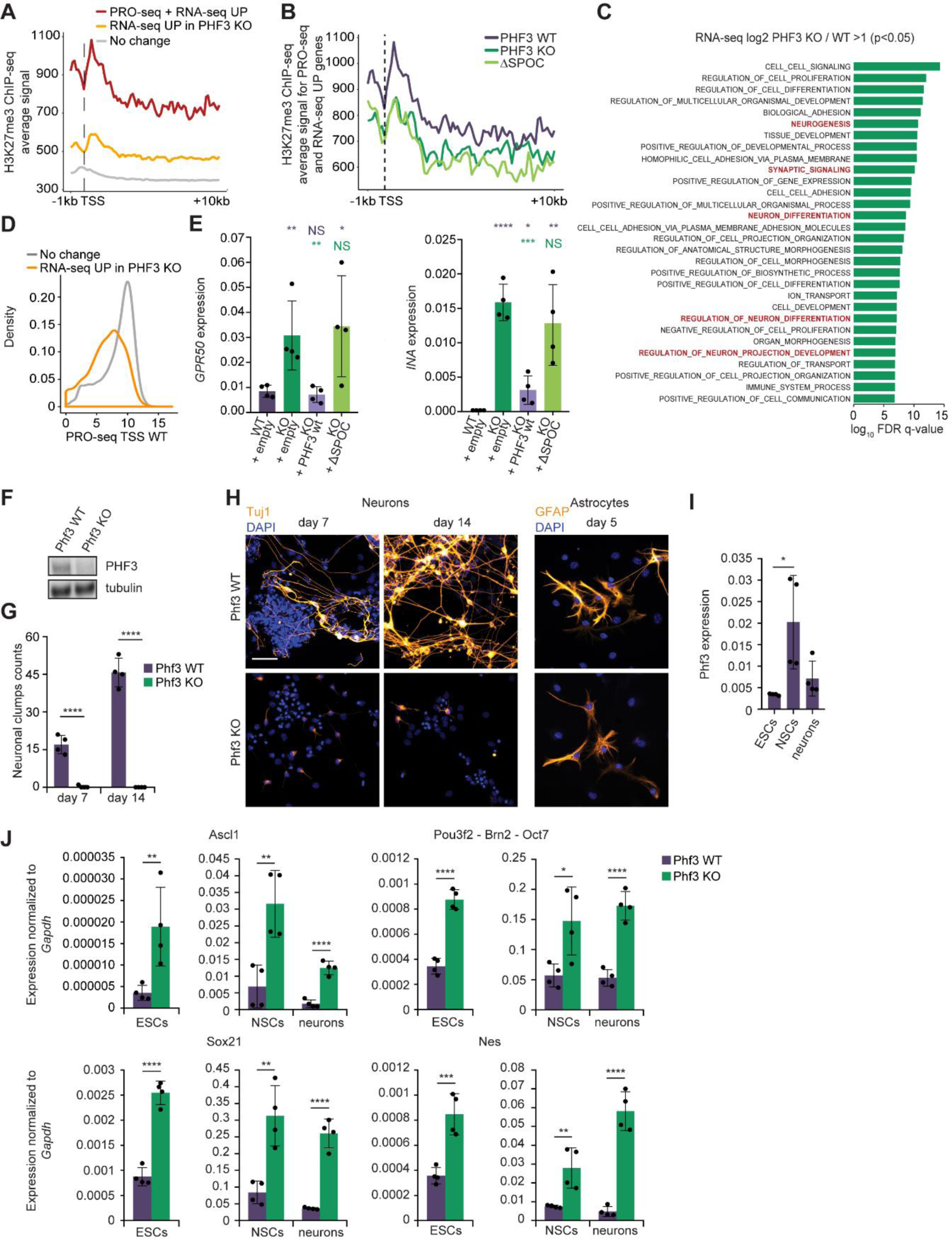
PHF3 negatively regulates neuronal gene expression and neuronal differentiation. (A) Composite analysis of H3K27me3 distribution and signal strength in PHF3 WT cells on TSS-gene body region for different gene categories based on RNA-seq and PRO-seq data. (B) Composite analysis of H3K27me3 distribution and signal strength in PHF3 WT, KO and ΔSPOC cells on TSS-gene body region for genes upregulated in RNA-seq and PRO-seq in PHF3 KO or ΔSPOC cells (fold change>2). (C) GO analysis of genes upregulated in PHF3 KO according to RNA-seq shows enrichment of genes involved in neurogenesis. GSEA ‘Biological processes’ tool was used. (D) Genes upregulated in PHF3 KO cells according to RNA-seq (fold change>2) have low expression levels in WT cells as judged by nascent transcription (PRO-seq) levels at TSS. (E) RT-qPCR analysis of INA and GPR50 expression levels in PHF3 WT and PHF3 KO with stable integration of mCherry empty vector, and KO-complemented cell lines stably expressing mCherry-PHF3 wild-type or ΔSPOC (mean±sd; n=4). The bars represent average expression from different clones as biological replicates. A t-test was performed by comparing expression levels with WT (violet asterisk) and KO (green asterisk). (F) CRISPR/Cas9 Phf3 knock-out in mESCs shows complete loss of protein by Western blotting. (G) Quantification of beta III tubulin (TuJ1)-positive neuronal clump formation in Phf3 WT and KO cells after 7 or 14 days of neuronal differentiation. (H) Representative immunofluorescence images of TuJ1-stained neurons and glial fibrillary acidic protein (GFAP)-stained astrocytes. Scale bar=40 μm. (I) Phf3 expression levels in embryonic stem cells (ESCs), neural stem cells (NSCs) and neurons determined by RT-qPCR (n=4). (J) Comparison of expression levels of different neuronal markers between Phf3 WT and KO ESCs, NSCs and neurons by RT-qPCR (n=4).

Our data define PHF3 as an elongation factor that negatively regulates mRNA stability, and thereby couples transcription, RNA processing and RNA metabolism.

### PHF3 negatively regulates neuronal genes

Genes with highly elevated transcripts in PHF3 KO and ΔSPOC are transcribed at low levels in WT cells and enriched for both the ‘repressive’ H3K27me3 mark and the ‘active’ H3K4me3, as well as pS5 (Figure 6A,D, S11). Genes that showed upregulation of both nascent and mature transcripts in PHF3 WT and ΔSPOC cells displayed a decrease in H3K27me3 (Figure 6B), suggesting that PHF3-mediated transcriptional derepression is coupled with the loss of Polycomb-mediated silencing. Overall, these data suggest that Pol II is in a ‘poised’ state and possibly undergoing fast turnover at this group of genes in WT cells (Ferrai et al., 2017; Price, 2018).

GO analysis of the upregulated mature transcripts in PHF3 KO and ΔSPOC HEK293T cells revealed an enrichment of neuronal genes (Figure 6C,E and Figure S6C,D), whereas downregulated or unaffected genes did not show any particular functional enrichment. For example, the neuronal genes *INA* and *GPR50* exhibited low transcript levels in WT cells, whereas loss of PHF3 or the PHF3 SPOC domain resulted in pronounced increase in transcript levels (Figure 4E, 6E). Exogenous expression of PHF3 in PHF3 KO cells reduced the expression of *INA* and *GPR50* to levels that were comparable to WT cells (Figure 6E). Importantly, PHF3 ΔSPOC failed to rescue the PHF3 KO phenotype (Figure 6E).

### Phf3 is required for neuronal differentiation of mESCs

Gene expression during neuronal development is frequently regulated at the level of mRNA stability (Deschenes-Furry et al., 2006). Many of the neuronal genes that we found to be directly repressed by PHF3, including *INA* (Yuan et al., 2017) and *GPR50* (Khan et al., 2016), have been implicated in different aspects of neurodevelopment. To examine a potential role for PHF3 in proper neuronal differentiation, we used CRISPR/Cas9 to generate Phf3 KO mouse embryonic stem cells (mESCs) (Figure 6F and Figure S12A). We differentiated Phf3 KO and WT mESCs into neural stem cells (NSCs), and subsequently into neurons or astrocytes. As expected, NSCs derived from WT mESCs could differentiate into beta III tubulin (TuJ1)-positive neurons organized in extensively connected neuronal clumps (Figure 6G,H). WT NSCs could also differentiate into glial fibrillary acidic protein (GFAP)-positive astrocytes (Figure 6H). In contrast, although NSCs derived from Phf3 KO mESCs formed astrocytes comparable to WT NSCs, they failed to differentiate into properly shaped and connected neurons (Figure 6G,H). Additionally, we found that Phf3 transcript levels were elevated in WT NSCs relative to WT mESCs, suggesting that Phf3 expression is regulated during neuronal differentiation (Figure 6I). Our findings suggest that Phf3 is required for proper terminal differentiation of NSCs, specifically into the neuronal lineage.

We hypothesized that loss of Phf3 triggers derepression of specific genes that must be tightly regulated for efficient neuronal differentiation. To test this, we analyzed Phf3 KO mESCs by RNA-seq. Indeed, loss of Phf3 led to the upregulation of several factors that are important for neuronal fate specification, such as Ascl1, Pou3f2, Sox21 and Nestin, in mESCs, NSCs and neurons (Figure 6J). The pioneer proneural transcription factor Ascl1 must be tightly regulated for the development and proliferation of NSCs, as well as for the differentiation of progenitors along the neuronal lineage (Vasconcelos and Castro, 2014). The transcription factor Pou3f2 (also called Oct7 or Brn2) acts downstream of Ascl1 and is required for the differentiation of neural progenitor cells into functional neurons (Wapinski et al., 2013). High levels of the intermediate filament Nestin are also expected to interfere with terminal neuronal differentiation (Lendahl et al., 1990). Upregulation of the transcription factor Sox21 induces premature expression of neuronal markers but also inhibits terminal neuronal differentiation (Ohba et al., 2004; Sandberg et al., 2005; Whittington et al., 2015). Additionally, the stemness marker Sox2, which promotes neural stemness specification and suppresses neuronal differentiation (Graham et al., 2003), was upregulated in Phf3 KO NSCs and neurons relative to WT controls (Figure S12D).

Whereas neuronal factors were upregulated upon loss of Phf3, the embryonic stemness markers Oct4 and Nanog showed reduced expression in Phf3 KO in NSCs and neurons relative to WT controls (Figure S12D). In line with the premature expression of neural markers, Phf3 KO mESCs showed accelerated exit from naïve pluripotency compared to WT mESCs, as observed by upregulation of Pou3f1 (Oct6), Fgf5, Otx2 and Pax6 within the first 24 h of differentiation (Figure S12E). Taken together, our data suggest that Phf3 KO mESCs fail to differentiate into neurons due to aberrant and precocious derepression of factors that regulate neuronal commitment and terminal differentiation.

## DISCUSSION

Here, we established PHF3 as a new Pol II regulator that couples transcription with mRNA stability. In addition, we discovered that PHF3 is required for proper neuronal differentiation by preventing the precocious expression of a subset of neuronal genes. We found that PHF3 binds to the Pol II CTD phosphorylated on S2 through a new CTD-binding domain called SPOC. While the Pol II CTD is the primary anchoring point for PHF3, this large mammalian protein can likely establish additional contacts with Pol II, such as through the TLD domain, which was shown to bind the jaw-lobe domain of Pol II in the case of the yeast homolog Bye1 (Kinkelin et al., 2013) (Figure S13). Bivalent interaction of PHF3 with Pol II, through the Pol II jaw-lobe domain and through the CTD, may explain the dual repressive function of PHF3 in transcription and mRNA stability.

How does PHF3 regulate transcription? We found that in the absence of PHF3, TFIIS is more strongly bound to the Pol II complex. PHF3 competes with TFIIS for binding to Pol II and thereby impedes the rescue of backtracked Pol II. Pol II backtracking is a widespread phenomenon and backtracked Pol II may be particularly susceptible to premature termination and fast turnover (Sheridan et al., 2019), which would explain why PHF3-regulated genes are low-expressed with low Pol II occupancy, but have open promoters (Figure S11). Our data suggest that PHF3 represses these genes by competing with TFIIS to prevent the rescue of backtracked Pol II and promote premature termination. In the absence of PHF3, TFIIS binds more strongly to Pol II lobe-jaw domain and stimulates productive elongation (Figure 7). Ongoing structural work with the full-length PHF3 is expected to further clarify the competitive binding mechanism of PHF3 with respect to TFIIS.

**Figure 7.**
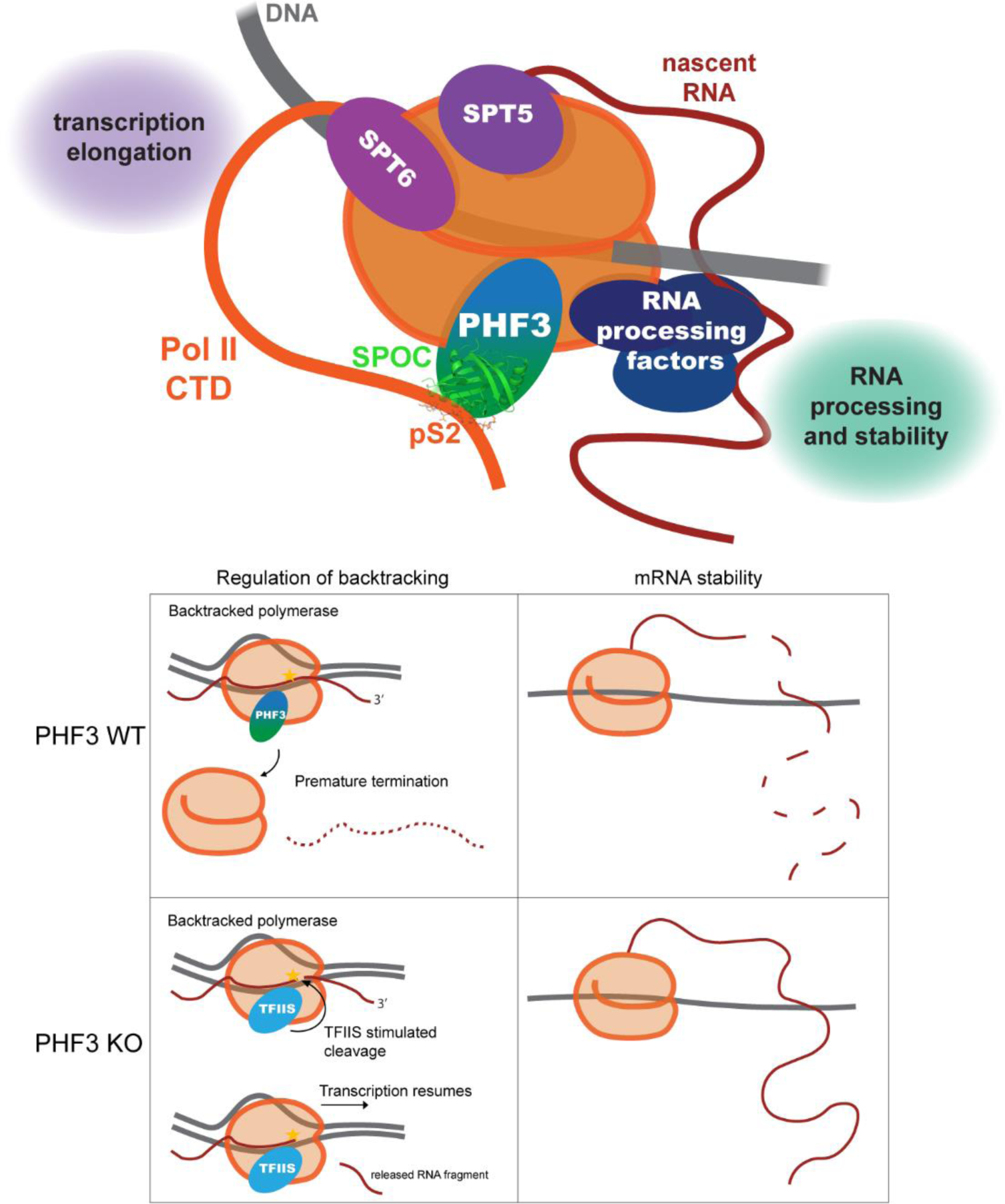
A model of PHF3-mediated gene repression. PHF3 binds Pol II CTD phosphorylated on Serine-2 and interacts with transcription elongation machinery and RNA processing factors. PHF3 represses transcription by competing with TFIIS and impeding Pol II rescue from backtracking. PHF3 is also a negative regulator of mRNA stability.

How does PHF3 regulate mRNA stability? PHF3 may regulate mRNA stability by modulating the recruitment of RNA-binding proteins (RBPs) that were identified in the PHF3 interactome and/or that associate with the Pol II CTD and regulate RNA processing. RNA processing factors could require PHF3 for binding to the CTD. Coupling between elongation and RNA processing through Pol II CTD becomes even more relevant considering recent findings that the CTD undergoes liquid-liquid phase separation (LLPS) (Boehning et al., 2018; Lu et al., 2018). LLPS of unphosphorylated CTD may facilitate Pol II clustering and transcriptional bursting during transcription initiation, whereas the subsequent LLPS wave of pS2-phosphorylated CTD may trigger Pol II clustering with RNA processing factors (Guo et al., 2019). PHF3 may promote the second wave of CTD LLPS through direct binding to pS2 CTD and RNA processing factors. While further work will elucidate how exactly PHF3 regulates mRNA stability, our results establish PHF3 as a new mammalian ‘synthegradase’ that coordinates transcription with mRNA decay (Bregman et al., 2011; Haimovich et al., 2013).

Why do genes react differently to PHF3 loss? Neuronal genes were enriched among 10% of de-repressed genes with high levels of both nascent and mature transcripts. These genes are low-expressed and Polycomb-repressed in WT cells but bear the marks of ‘poised’ Pol II (H3K4me3, Pol II pS5). Poised Pol II is found in ESCs as well as differentiating and post-mitotic cells (Ferrai et al., 2017). During neuronal differentiation, poised Pol II primes neuronal transcription factors for activation whilst keeping non-neuronal genes silenced (Ferrai et al., 2017). Poised Pol II may experience high levels of backtracking due to increased GC content, specific promoter elements or chromatin configuration (Core and Adelman, 2019; Gomez-Herreros et al., 2012); PHF3 would prevent efficient TFIIS-mediated rescue from backtracking and induce premature termination. Reactivation of these genes would thus be highly dependent on TFIIS, which may be the reason for their marked de-repression in PHF3 KO cells where TFIIS would gain more access to Pol II.

In differentiated HEK293T cells, PHF3 represses neuronal genes and thereby may contribute to the maintenance of non-neuronal cell identity. During neuronal differentiation of mESCs, Phf3 fine-tunes expression of neuronal genes to ensure their timely and adequate expression levels during differentiation. Phf3 KO mESCs fail to undergo neuronal differentiation, implying that PHF3 is required for neuronal development. Indeed, a Phf3 KO mouse generated by the International Mouse Phenotyping Consortium (IMPC) exhibits neuronal dysfunction in the form of impaired auditory brainstem response and impaired startle reflex (www.mousephenotype.org).

PHF3 function in the regulation of transcription and mRNA stability may be important beyond development. PHF3 was found to be significantly downregulated in glioblastoma – the most common undifferentiated brain tumour (Fischer et al., 2001). Interestingly, ASCL1, POU3F2, and SOX2, which were all derepressed in Phf3 KO cells, have been implicated in the maintenance and tumorigenicity of glioblastoma (Rheinbay et al., 2013; Suva et al., 2014). Our data suggest that PHF3 downregulation may drive glioblastoma via derepression of transcription factors that regulate neuronal differentiation.

## SUPPLEMENTAL INFORMATION

Supplemental information includes 13 figures.

## AUTHOR CONTRIBUTIONS

L.A. generated endogenous PHF3 GFP-tagged and SPOC-deleted cell lines, performed all ChIP and SLAM-seq experiments, purified SPOC and Pol II, performed and analyzed FCS experiments; V.F. designed and performed the analysis of all sequencing data and conceptually drove the project; M.B. generated PHF3 KO (with S.K.), performed and analyzed co-immunoprecipitation, EU incorporation, neuronal differentiation experiments, performed RNA-seq and PRO-seq (with U.S.), generated PHF3 KO cell lines stably expressing PHF3 and ΔSPOC PHF3 and performed complementation experiments;

I.G. solved the SPOC structures; A.K. performed and analyzed FA experiments; U.S. performed PRO-seq experiments; M.P. supervised and analyzed FCS experiments; S.K. purified SPOC; E.B. analyzed mass spectrometry data; K.M. supervised mass spectrometry analysis; G.L. and A.V. analyzed HEK293 RNA-seq data; M.L. supervised ESC genome-engineering and differentiation experiments; R.P. supervised initial Pol II ChIP experiments; A.S. supervised the analysis of HEK293 RNA-seq data; A.A. designed and supervised the analysis of sequencing data; R.S. supervised FA experiments; C.B. supervised Pol II *in vitro* assays; K.D.C. supervised X-ray analysis; D.S. conceived the study, performed, supervised, analyzed experiments and wrote the manuscript.

## ACKNOWLEDGEMENTS

D.S. thanks Claudine Kraft, Renée Schroeder, Verena Jantsch, Franz Klein and Peter Schlögelhofer for support. We thank Anita Testa Salmazo for help with purifying Pol II; Matthias Geyer and Robert Düster for sharing DYRK1A kinase; Goran Kokic for design of the arrest assay sequences; Petra van der Lelij for help with generating mESC KO; Stefan Ameres, Nina Fasching and Brian Reichholf for advice on SLAM-seq and for sharing reagents; Krzysztof Chylinski for advice regarding CRISPR/Cas9 methodology; VBCF Protein Technologies facility for purifying PHF3 and providing gRNAs and Cas9; VBCF NGS facility for sequencing; Monoclonal antibody facility at the Helmholtz center for Pol II antibodies; Friedrich Propst and Elzbieta Kowalska for advice and for sharing materials; Egon Ogris for sharing materials; Martin Eilers for recommending a ChIP-grade TFIIS antibody; Susanne Opravil, Otto Hudecz, Markus Hartl and Natascha Hartl for mass spectrometry analysis; staff of the X-ray beamlines at the ESRF in Grenoble for their excellent support; Christa Bücker, Anton Meinhart, Clemens Plaschka and members of Slade lab for critical comments on the manuscript; Life Science Editors for editing assistance. M.B. and D.S. acknowledge support by the FWF-funded DK ‘Chromosome Dynamics’. U.S. is supported by the L’Oreal for Women in Science Austria Fellowship and the Austrian Science Fund (FWF T 795-B30). M.L is supported by the Vienna Science and Technology Fund (WWTF, VRG14-006). R.S. is supported by the Czech Science Foundation (15-17670S), Ministry of Education, Youths and Sports of the Czech Republic (CEITEC 2020 project (LQ1601)), and the European Research Council (ERC) under the European Union’s Horizon 2020 research and innovation programme (grant agreement No. 649030); this publication reflects only the author’s view and the Research Executive Agency is not responsible for any use that may be made of the information it contains. K.D.C. research is supported by the Austrian Science Fund (FWF) Projects I525 and I1593, P22276, P19060 and W1221, Federal Ministry of Economy, Family and Youth through the initiative ‘Laura Bassi Centres of Expertise’, funding from the Centre of Optimized Structural Studies N°253275, the Wellcome Trust Collaborative Award (201543/Z/16), COST action BM1405 Non-globular proteins - from sequence to structure, function and application in molecular physiopathology (NGP-NET), the Vienna Science and Technology Fund (WWTF LS17-008), and by the University of Vienna. This project was funded by the MFPL start-up grant, the Vienna Science and Technology Fund (WWTF LS14-001) and the Austrian Science Fund (P31546-B28 and W1258 “DK: Integrative Structural Biology”) to D.S.

## KEY RESOURCES TABLE

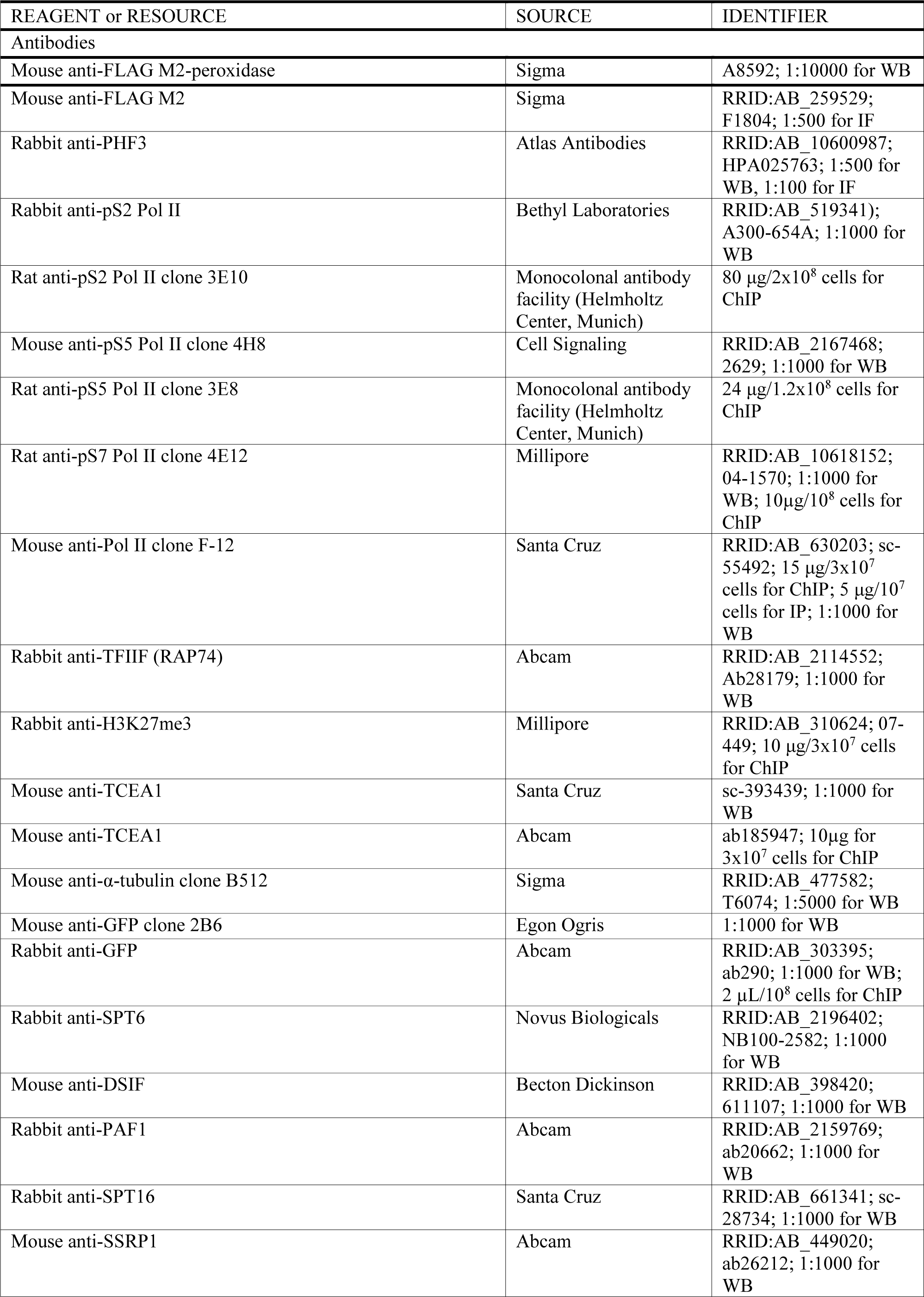
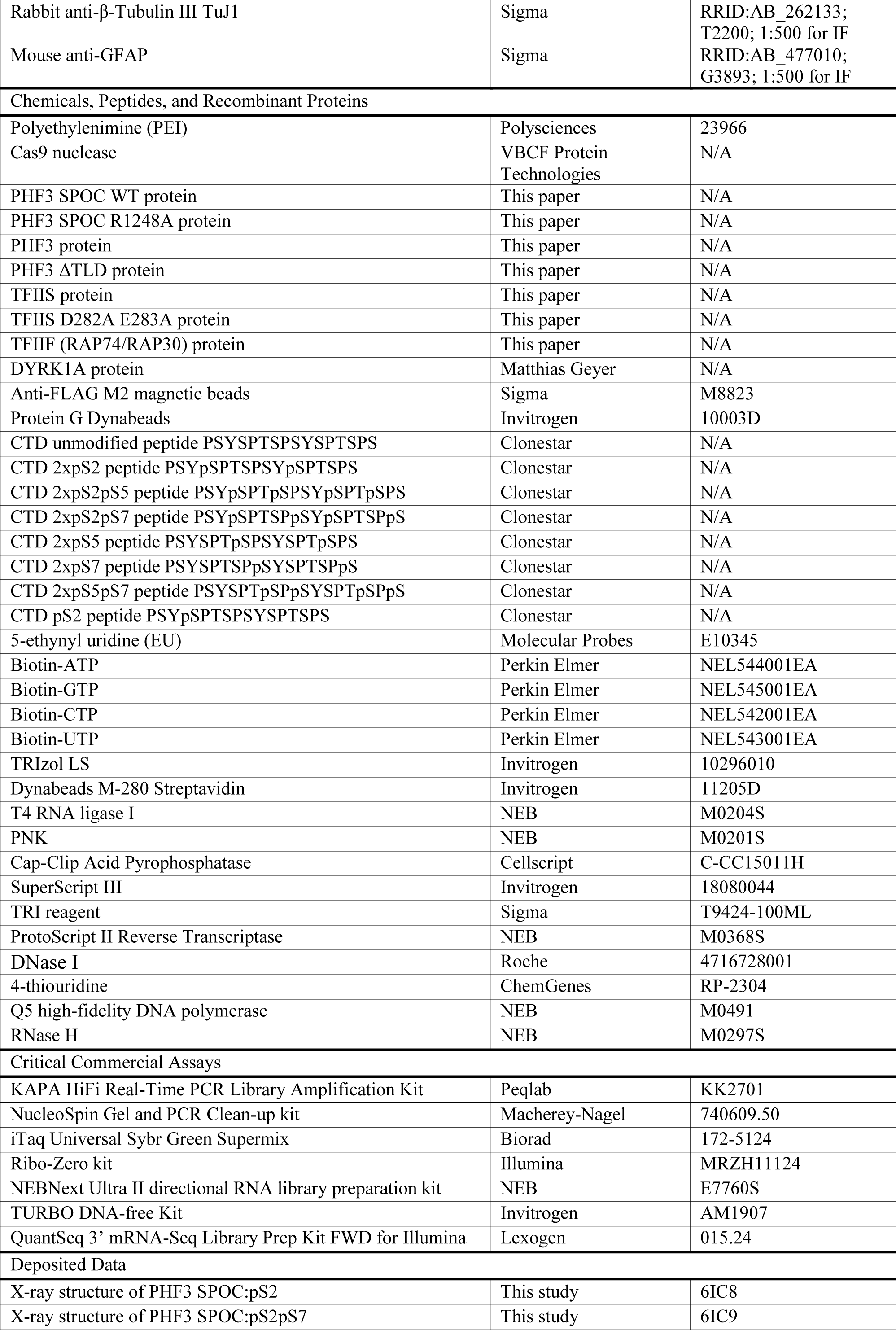
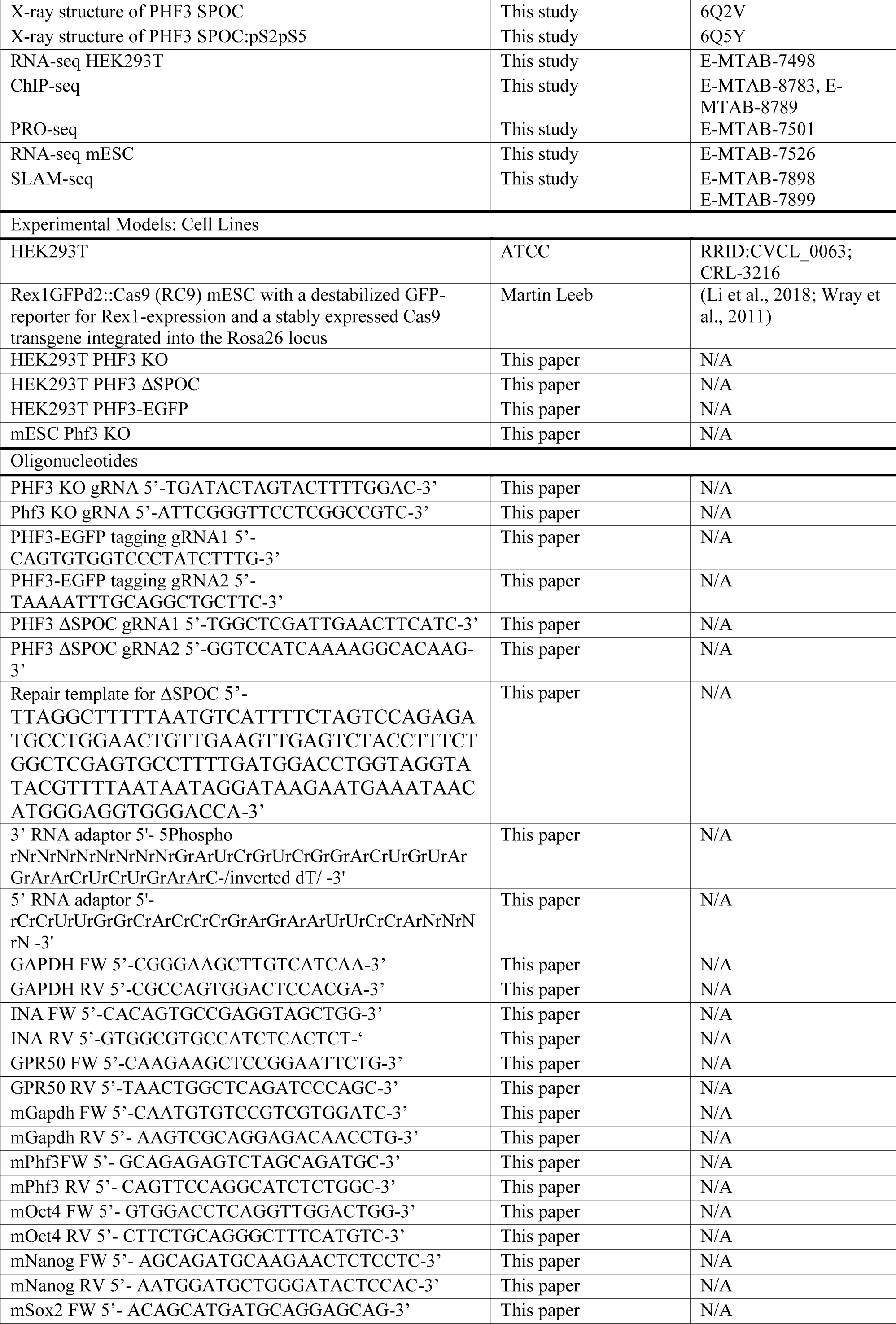
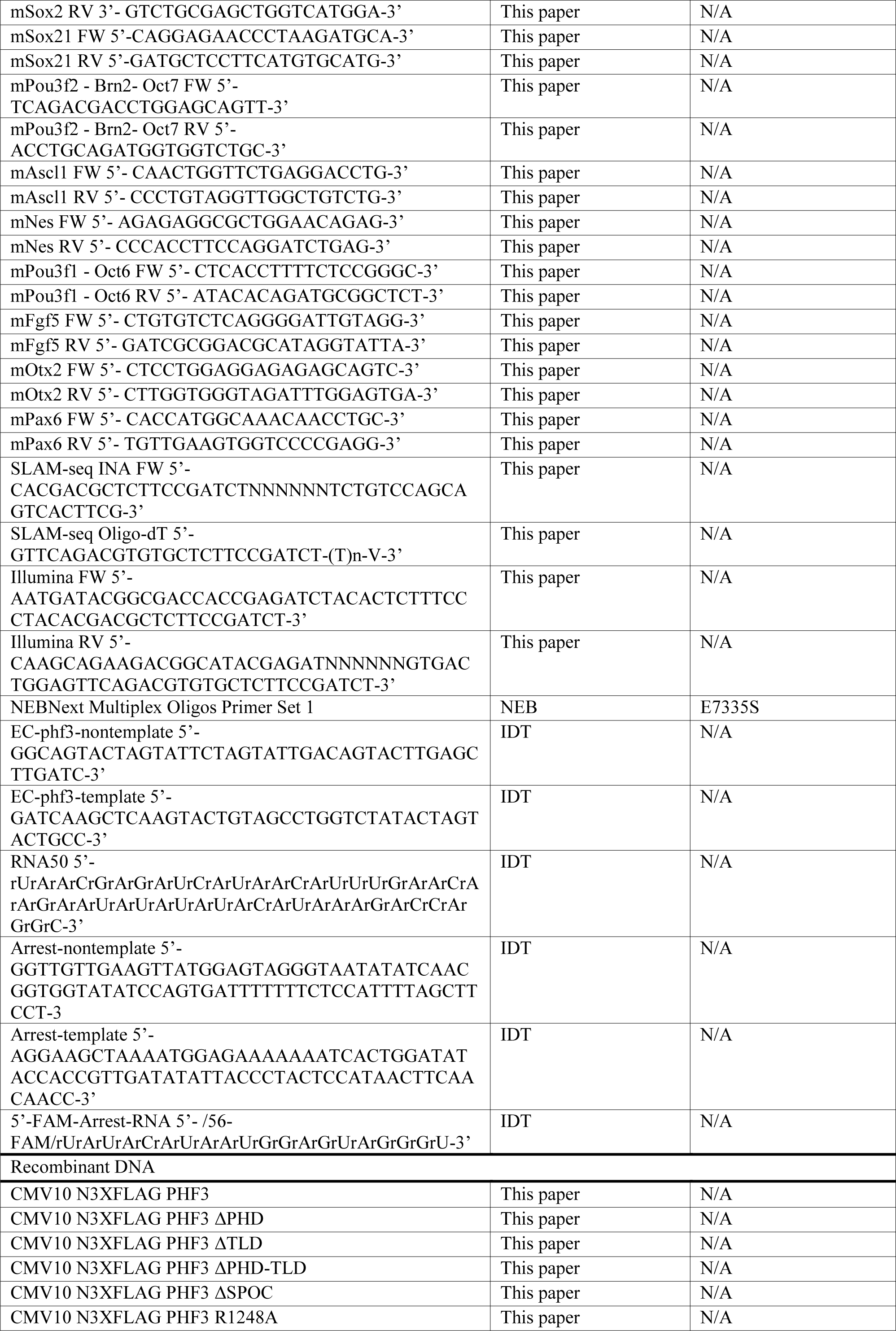
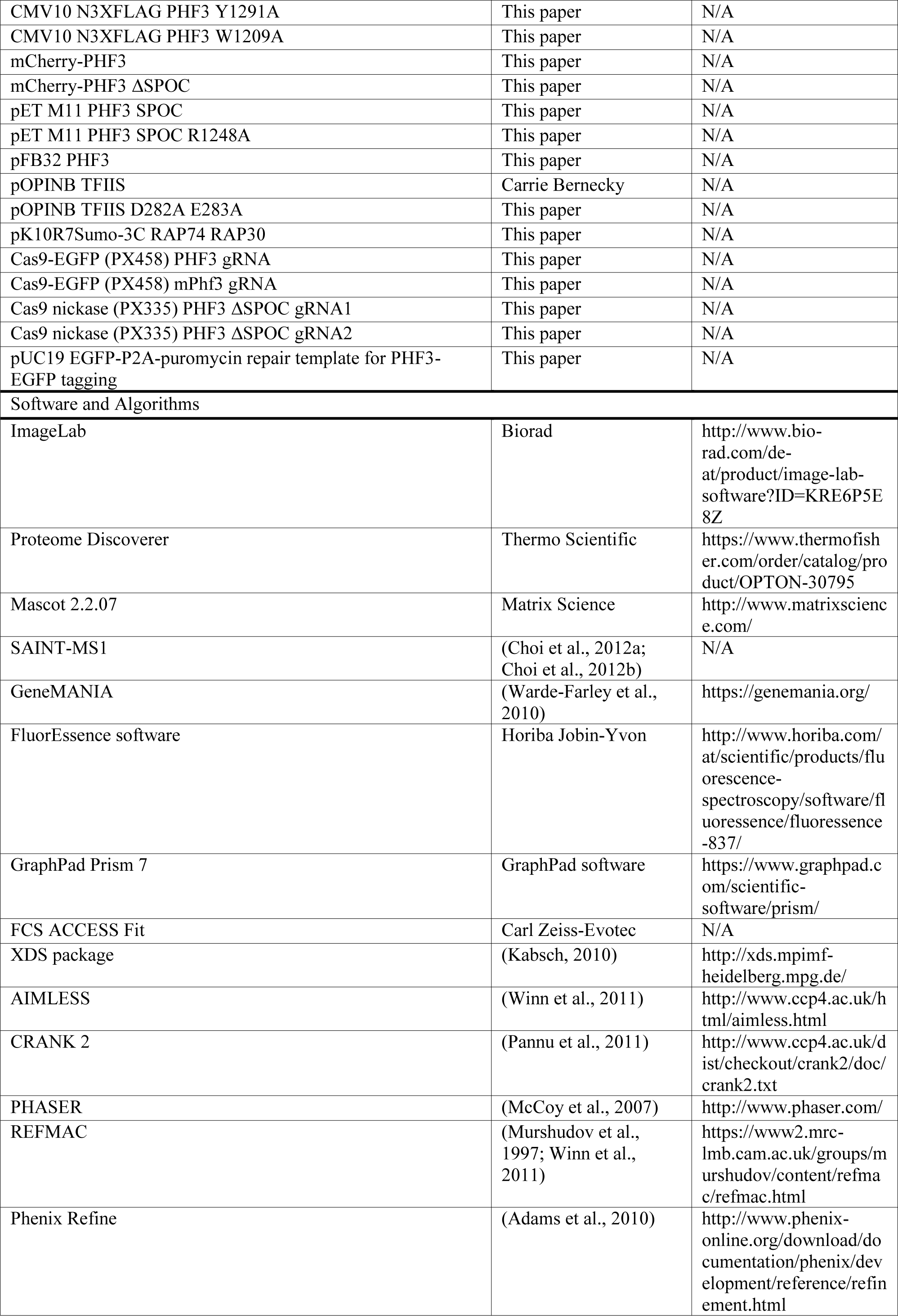
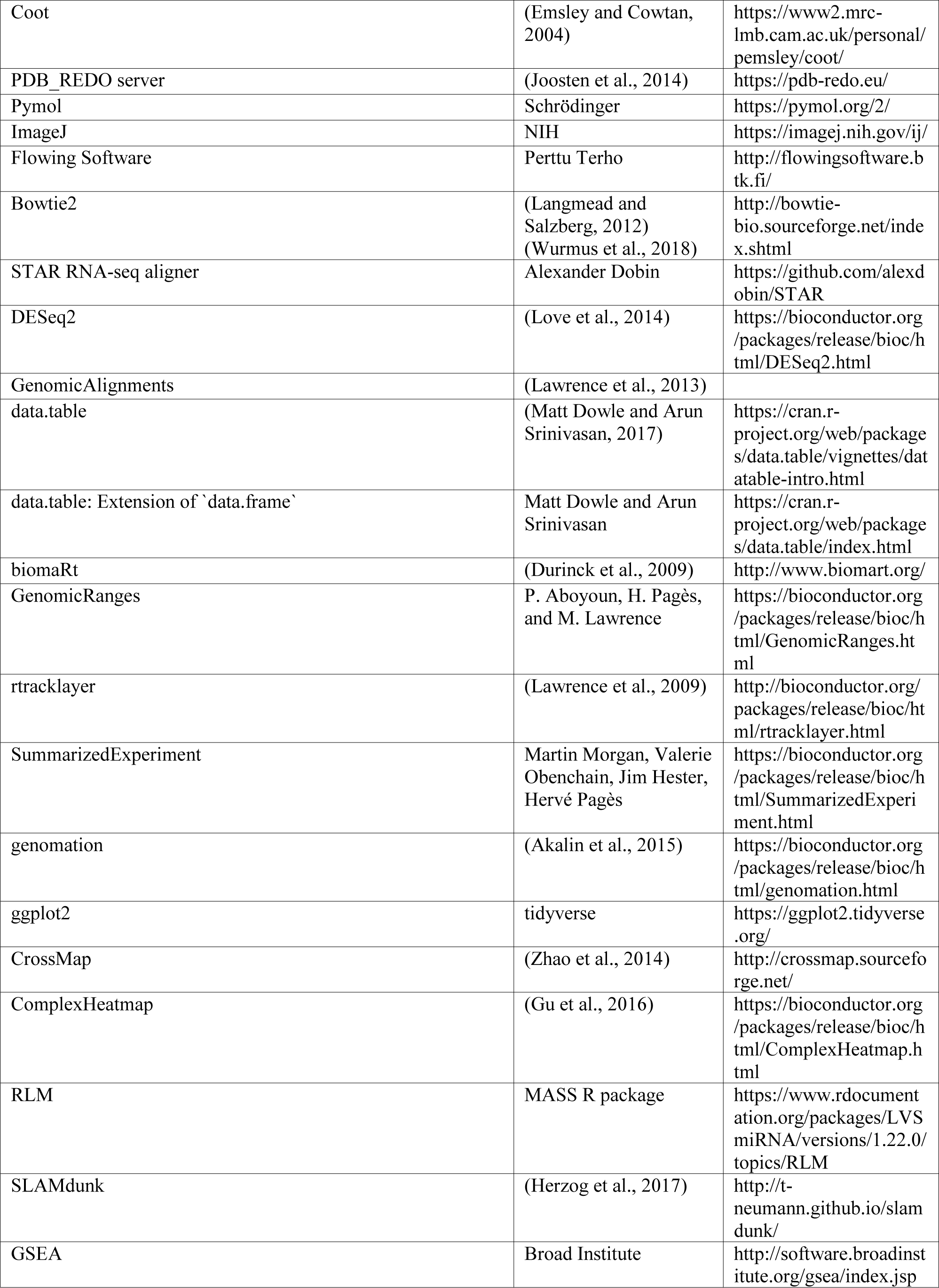
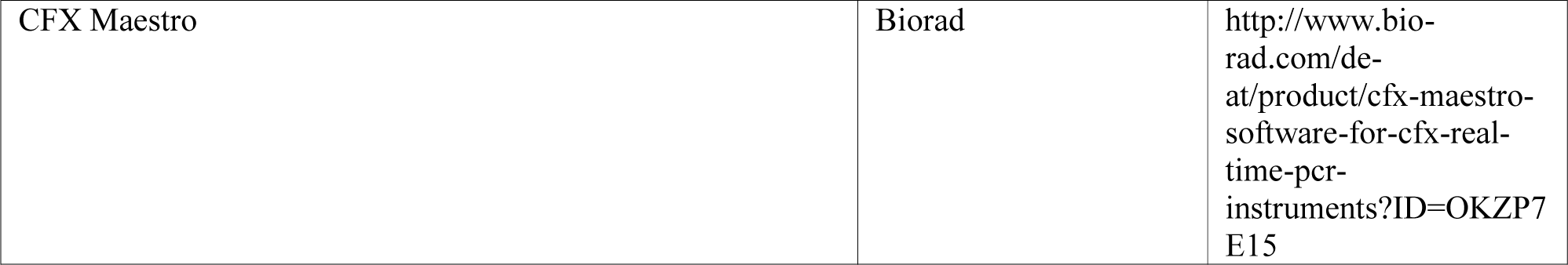

## CONTACT FOR REAGENT AND RESOURCE SHARING

Further information and requests for reagents may be directed to and will be fulfilled by the Lead Contact, Dr. Dea Slade, University of Vienna.

## EXPERIMENTAL MODEL AND SUBJECT DETAILS

### Cell lines and cell culture

HEK293T cells were grown in Dulbecco’s Modified Eagle’s Medium (DMEM 4.5 g/L glucose) (Sigma) supplemented with 10% fetal bovine serum (Sigma), 1% L-glutamine (Sigma), 1% penicillin-streptomycin (Sigma) under 5% CO_2_ at 37°C. mESCs were cultured on 0.2% gelatin coated plates in ES-DMEM medium supplemented with LIF and 2i as described previously (Leeb et al., 2015). To generate CRISPR/Cas9 PHF3 KO, gRNA targeting exon 3 was cloned between BbsI sites under the U6 promoter in the plasmid encoding Cas9-EGFP (pX458) (Ran et al., 2013). gRNA sequence for human PHF3 was 5’-TGATACTAGTACTTTTGGAC-3’ and for mouse Phf3 5’-ATTCGGGTTCCTCGGCCGTC-3’. 48h after transfection with polyethylenimine (PEI; Polysciences), GFP-positive HEK293T cells were FACS-sorted and allowed to recover in culture for 4 to 7 days. Cells were subsequently FACS-sorted and GFP-negative cells were seeded 1 cell/well in 96-wells plates. After 14 to 20 days, surviving clones were expanded in culture, genomic DNA was isolated and PCR-amplified Cas9 target region was sequenced. To generate CRISPR/Cas9 Phf3 KO in mESCs, Rex1GFPd2::Cas9 (RC9) ES cells were used that carry a destabilized GFP-reporter for Rex1-expression and a stably expressed Cas9 transgene integrated into the Rosa26 locus. ES cells were co-transfected with 0.5 µg gRNA-expressing plasmid and 0.1 µg of dsRed expressing plasmid. After 2 days dsRed-positive cells were FACS-sorted and plated at clonal density into 60 mm TC-dishes. After 7 days colonies were picked and expanded in 96-well plates. To identify KO clones, genomic DNA was isolated and PCR-amplified Cas9 target region was sequenced. To generate CRISPR/Cas9 endogenously GFP-tagged PHF3, two gRNAs targeting 3’ PHF3 terminus were designed (5’-CAGTGTGGTCCCTATCTTTG-3’ and 5’-TAAAATTTGCAGGCTGCTTC-3’) and cloned into the plasmid pX335 encoding Cas9 nickase (Cong et al., 2013). Plasmid-borne repair template consisted of EGFP-P2A-puromycin flanked by 1.5 kb sequences homologous to the targeted genomic region. HEK293T cells were transfected with 2 µg of each of the plasmids encoding Cas9 nickase and one of the two gRNAs and 4 µg of the repair template. Two weeks after transfection, GFP positive cells were sorted by FACS. Two days after the sorting 0.5 µg/mL puromycin was added to the culture medium. After 1-2 weeks, surviving colonies were picked and expanded, genomic DNA was extracted and positive clones were identified by PCR. To generate CRISPR/Cas9 PHF3 ΔSPOC, one gRNA target site on either side of the SPOC domain was selected in such a way that the PAMs are within the deleted fragment (5’-TGGCTCGATTGAACTTCATC-3’ and 5’-GGTCCATCAAAAGGCACAAG-3’). A 150 bp ssDNA repair template was designed which introduces an XhoI restriction site at the junction of the two breaks without shifting the reading frame or altering the amino acid sequence (5’-TTAGGCTTTTTAATGTCATTTTCTAGTCCAGAGATGCCTGGAACTGTTGAAGTTGAGTCTA CCTTTCTGGCTCGAGTGCCTTTTGATGGACCTGGTAGGTATACGTTTTAATAATAGGATAAGAATGAAATAACATGGGAGGTGGGACCA-3’). S-phase synchronized HEK293T cells were electroporated with 10 µg of purified Cas9, 12 µg of each *in vitro* transcribed gRNA and 4 µM repair template. Genomic DNA from the clones was PCR-amplified to check for the deletion of 4.5 kb and digested with XhoI to ensure that the repair template was used for homologous recombination-mediated repair. To generate stable cell lines expressing mCherry-PHF3 constructs, HEK293T PHF3 KO cells were transfected with 2 μg of plasmid and PEI in 6-well plates. After 48 h, half of the cells were transferred to 10 cm dishes and grown in medium supplemented with 0.25 µg/mL of puromycin. After 2-3 weeks, surviving colonies were picked using glass cylinders and monoclonal populations were expanded in culture. Positive clones were validated by Western blot.

## METHOD DETAILS

### Constructs

Human PHF3 was amplified from HEK293T cDNA and cloned into CMV10 N3XFLAG (Sigma) between NotI and XbaI. PHF3 truncation constructs were generated by tripartite ligation of BsaI-flanking fragments according to the Golden Gate cloning principle (Engler et al., 2009). PHF3 and ΔSPOC constructs used for complementation were cloned into mCherry IRES puromycin vector (Clontech) between AgeI and NotI. Site-directed mutagenesis was performed according to the FastCloning protocol (Li et al., 2011). For bacterial expression, PHF3 SPOC domain (1199-1356aa) was cloned into pET M11 between NcoI and XhoI for N-terminal His_6_ fusion. For insect cell expression, PHF3 was cloned into pFB32 for N-terminal His_6_ fusion and C-terminal Strep fusion. TFIIS mutant (TFIIS^M^) was generated by site-directed mutagenesis to introduce D282A E283A mutations in pOPINB TFIIS for bacterial expression. pK10R7Sumo-3C TFIIF was generated by amplifying RAP74 and RAP30 from HEK293T cDNA; RAP74 was cloned between BamHI and NotI for N-terminal SUMO-His_10_ fusion, RAP30 was cloned between NdeI and KpnI.

### Protein purification

SPOC was expressed in *E. coli* Rosetta2 (DE3) cells (Novagen) and purified by affinity chromatography using HisTrap HP column (GE Healthcare) equilibrated in 25 mM Tris pH 7.4, 500 mM NaCl, 20 mM imidazole, followed by TEV cleavage of the His_6_ tag and size exclusion chromatography using Sephacryl S-200 (GE Healthcare) equilibrated in 25 mM Tris pH 7.4, 25 mM NaCl and 1 mM DTT. TFIIS or TFIIS^M^ were expressed in *E. coli* Rosetta2 (DE3) cells (Novagen) and purified by affinity chromatography using HisTrap HP column (GE Healthcare) equilibrated in 25 mM Tris pH 7.4, 500 mM NaCl, 20 mM imidazole, and by size exclusion chromatography using Sephacryl S-200 (GE Healthcare) equilibrated in 5 mM Hepes pH 7.25, 100 mM NaCl, 10 µM ZnCl_2_ and 10 mM DTT. TFIIF (RAP74/RAP30) was expressed in *E. coli* Rosetta2 (DE3) cells (Novagen) and purified by affinity chromatography using HisTrap HP column (Ge Healthcare) equilibrated in 25 mM Tris pH 7.4, 500 mM NaCl, 20 mM imidazole, followed by 3C cleavage of the SUMO-His_10_ tag, cation exchange chromatography using HiTrap SP column (GE Healthcare) equilibrated in 50 mM Hepes pH 7, 150 mM KCl, 10% glycerol and 2 mM DTT, and by size exclusion chromatography using Sephacryl S-200 (GE Healthcare) equilibrated in 20 mM Hepes pH 7.5, 100 mM NaCl, 10% glycerol and 2 mM DTT. PHF3 was expressed from the EMBacY bacmid in Sf9 cells with an N-terminal His-tag and a C-terminal Strep-tag. PHF3 was purified by affinity chromatography using HisTrap FF column (GE Healthcare) equilibrated in 50 mM Tris-Cl pH 8, 300 mM NaCl, 0.5 mM TCEP, 20 mM imidazole, followed by anion exchange chromatography using HiTrap ANX (high sub) FF column (GE Healthcare) equilibrated in 50 mM Tris-Cl pH 8, 100 mM NaCl, 0.5 mM TCEP, 10% glycerol, and size exclusion chromatography using Superose 6 (GE Healthcare) equilibrated in 50 mM Tris pH 8.0, 300 mM NaCl, 0.5 mM TCEP. Pol II was purified from pig thymus as previously described (Bernecky et al., 2016).

### Immunoprecipitation

For immunoprecipitation of exogenously expressed FLAG-PHF3 constructs, a 10 cm dish of transfected HEK293T cells was used. Cells were harvested 48 h after transfection and lysed in lysis buffer (50 mM Tris-Cl pH 8, 150 mM NaCl, 1% Triton, 1x protease inhibitors, 2 mM Na_3_VO_4_, 1 mM PMSF, 2 mM NaF, 50 units/mL benzonase and 1 mM DTT) for 1 h at 4°C. 10% of the cleared lysate was kept as input and the rest was incubated for 2 h on a rotating wheel at 4°C with anti-FLAG M2 magnetic beads (Sigma). For Pol II IP, Protein G Dynabeads (Invitrogen) were washed twice with TBS and incubated with 5 μg of mouse anti-FLAG M2 (Sigma), rabbit anti-pS2 Pol II (Abcam ab5095), mouse anti-pS5 Pol II 4H8 (Abcam ab5408), rat anti-pS7 Pol II clone 4E12 (Millipore) or mouse anti-Pol II clone F-12 (Santa Cruz) antibodies for 1 h on a rotating wheel at room temperature. Beads were washed twice with TBS and cleared lysates were added for immunoprecipitation on a rotating wheel at 4°C ON. For immunoprecipitation of endogenously tagged PHF3-GFP, two 15 cm dishes were used for each cell line. Cells were harvested and lysed in lysis buffer (as above but without DTT). The lysates were incubated on a rotating wheel at 4°C ON with rabbit anti-GFP (Abcam ab290) antibody. The samples were added to protein G Dynabeads (Invitrogen) and incubated on a rotating wheel at 4°C for 6 h. Beads were subsequently washed three times with TBS and immunoprecipitated proteins were eluted twice with 0.1 M glycine pH 2 and neutralized with Tris-Cl pH 9.2. For Western blot, 2% of the input and 20% of the eluate were loaded for each sample. For mass spectrometry analysis of FLAG-PHF3 or Pol II interactome, immunoprecipitations were performed as described above and samples were processed for on beads digestion.

### Mass spectrometry

Beads were eluted three times with 20 µL 100 mM glycine and the combined eluates were adjusted to pH 8 using 1 M Tris-HCl pH 8. Disulfide bonds were reduced with 10 mM DTT for 30 min before adding 25 mM iodoacetamide and incubating for another 30 min at room temperature in the dark. Remaining iodoacetamide was quenched by adding 5 mM DTT and the proteins were digested with 300 ng trypsin (Trypsin Gold, Promega) overnight at 37°C. The digest was stopped by addition of 1% trifluoroacetic acid (TFA), and the peptides were desalted using C18 Stagetips. NanoLC-MS analysis was performed using the UltiMate 3000 HPLC RSLC nano system (Thermo Scientific) coupled to a Q Exactive mass spectrometer (Thermo Scientific), equipped with a Proxeon nanospray source (Thermo Scientific). For FLAG-PHF3 IP samples, peptides were loaded onto a trap column (PepMap C18, 5 mm × 300 μm ID, 5 μm particles, 100 Å pore size; Thermo Scientific) followed by the analytical column (PepMap C18, 500 mm × 75 μm ID, 3 μm, 100 Å; Thermo Scientific). The elution gradient started with the mobile phases: 98% A (water/formic acid, 99.9/0.1, v/v) and 2% B (water/acetonitrile/formic acid, 19.92/80/0.08, v/v/v), increased to 35% B over the next 120 min followed by a 5-min gradient to 90% B, stayed there for five min and decreased in 5 min back to the gradient 98% A and 2% B for equilibration at 30°C. The Q Exactive mass spectrometer was operated in data-dependent mode, using a full scan followed by MS/MS scans of the 12 most abundant ions. For peptide identification, the .RAW-files were loaded into Proteome Discoverer (version 1.4.0.288, Thermo Scientific). The resultant MS/MS spectra were searched using Mascot 2.2.07 (Matrix Science) against the Swissprot protein sequence database, using the taxonomy human. The peptide mass tolerance was set to ±5 ppm and the fragment mass tolerance to ± 0.03 Da. The maximal number of missed cleavages was set to 2. The result was filtered to 1% FDR using Percolator algorithm integrated in Proteome Discoverer (Elias and Gygi, 2007). For Pol II IP samples, a pre-column for sample loading (Acclaim PepMap C18, 2 cm × 0.1 mm, 5 μm, Thermo Scientific), and a C18 analytical column (Acclaim PepMap C18, 50 cm × 0.75 mm, 2 μm, Thermo Scientific) were used, applying a segmented linear gradient from 2% to 35% and finally 80% solvent B (80% acetonitrile, 0.1% formic acid; solvent A 0.1% formic acid) at a flow rate of 230 nL/min over 120 min. Eluting peptides were analyzed on a Q Exactive HF-X Orbitrap mass spectrometer (Thermo Scientific), which was coupled to the column with a customized nano-spray EASY-Spray ion-source (Thermo Scientific) using coated emitter tips (New Objective). The mass spectrometer was operated in data-dependent acquisition mode (DDA), survey scans were obtained in a mass range of 375-1500 m/z with lock mass activated, at a resolution of 120k at 200 m/z and an AGC target value of 3E6. The 8 most intense ions were selected with an isolation width of 1.6 m/z, fragmented in the HCD cell at 27% collision energy and the spectra recorded for max. 250 ms at a target value of 1E5 and a resolution of 30k. Peptides with a charge of +1 or >+6 were excluded from fragmentation, the peptide match feature was set to preferred, the exclude isotope feature was enabled, and selected precursors were dynamically excluded from repeated sampling for 20 seconds. Raw data were processed using the MaxQuant software package (version 1.6.0.16; (Tyanova et al., 2016)) and the Uniprot human reference proteome (July 2018, www.uniprot.org) as well as a database of most common contaminants. The search was performed with full trypsin specificity and a maximum of three missed cleavages at a protein and peptide spectrum match false discovery rate of 1%. Carbamidomethylation of cysteine residues were set as fixed, oxidation of methionine, phosphorylation of serine, threonine and tyrosine, and N-terminal acetylation as variable modifications.

### Analysis of mass spectrometry data

For the analysis of the PHF3 interactome (FLAG-PHF3 IP), SAINT-MS1 was used as a statistical tool to determine the probability of protein-protein interactions (Choi et al., 2012a). Prior to analysis with SAINT-MS1 (Choi et al., 2012a) the label-free quantification data were cleaned by removing bait and common laboratory contaminants (Choi et al., 2012b). The controls (empty vector) was used simultaneously to estimate the parameters of the false interaction probability distributions. SAINT-MS1 was run for each method and fraction separately with 5000 and 10000 burn-in and sampling iterations, respectively. Protein areas were normalized to obtain a median protein ratio of one between samples. Fold changes were calculated based on these normalized protein areas. For the analysis of differential Pol II interactome between PHF3 WT and KO cells, label-free quantification the “match between runs” feature and the LFQ function were activated in the MaxQuant software package (Tyanova et al., 2016). Downstream data analysis was performed using the LFQ values in Perseus (version 1.6.2.3; (Tyanova et al., 2016)). Mean LFQ intensities of biological replicate samples were calculated and proteins were filtered for at least two quantified values being present in the three biological replicates. Missing values were replaced with values randomly selected from a normal distribution (with a width of 0.3 and a median downshift of 1.8 standard deviations of the sample population). To determine differentially enriched proteins we used the LIMMA package in R (version 3.5.1.) and applied the Benjamini-Hochberg correction for multiple testing to generate adjusted p-values.

### Fluorescence Anisotropy (FA)

All measurements were conducted on a FluoroMax-4 spectrofluorometer (Horiba Jobin-Yvon). The instrument was equipped with a thermostated cell holder with a Neslab RTE7 water bath (Thermo Scientific). The system was operated by FluorEssence software (version 2.5.3.0 and V3.5, Horiba Jobin-Yvon). All measurements were performed at 20^°^C in 25 mM Tris pH 8, 25 mM NaCl, 1 mM DTT. CTD peptides were labelled N-terminally with 5,6-carboxyfluorescein (FAM λex=467 nm, λem=517 nm; Clonestar). 10 nM CTD peptide (in a volume of 1.4 ml) was titrated with increasing amounts of SPOC protein. Each data point is an average of three measurements. The binding isotherms were generated by non-liner regression analyses with the software package GraphPad Prism 7 (GraphPad Software, La Jolla).

### Fluorescence Correlation Spectroscopy (FCS)

0.2 mg of lyophilized pS2 and pS2pS5 peptides (Clonestar) were dissolved in 30 µl DMSO (Sigma-Aldrich). One equivalent of Atto-488 NHS ester (Atto-tec GmbH) was added to three equivalents of peptide in DMSO followed by five equivalents of DIPEA (Sigma-Aldrich) and incubated for 16 h at room temperature protected from light. The reaction mixture was diluted to 50% (v/v) DMSO with water and purified by reverse-phase HPLC on a C18 column (Agilent Technologies) using a 10-70% gradient of acetonitrile + 0.1% TFA to water + 0.1% TFA over 16 min, followed by flushing at 90% acetonitrile + 0.1% TFA for 2 min. The desired fractions were lyophilized and stored at -20°C. A ConfoCor 2 spectrofluorimeter (Carl Zeiss-Evotec) equipped with an air-cooled Argon-laser (LASOS Lasertech GmbH; intensity 70 μW) and a water immersion objective (C-Apochromat 63 ×/ 1.2 W Corr) was used for monitoring changes in diffusion behavior due to binding of 15 nM labeled pS2 or pS2pS5 peptide to dilution series of 40 to 0.2 µM of SPOC. All FCS measurements were performed in a 1536-well glassplate (Greiner Bio-One) and a sample volume of 5 µL at 20°C. The diameter of the pinhole was set to 35 μm and the confocal volume was calibrated using 4 nM of Atto 488 dye (*D*_trans_ = 4.0 × 10^-10^ m^2^ s^−1^). Intensity fluctuations were recorded by an avalanche photodiode (SPCM-CD 3017) in photon counting mode over a time period of 15 sec and repeated 8 times for each sample. The normalized autocorrelation function *G*(τ) describes the observed fluctuations of the fluorescence intensity *δF*(*t*) from the mean intensity at any time compared to fluctuations at time *t* + *τ*. It is given by 

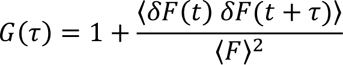

where the angular brackets represent the ensemble average, 〈F〉 denotes the mean intensity, and τ is known as the delay or correlation time interval over which the fluctuations are compared. For a single diffusing species of Brownian motion in a 3D Gaussian confocal volume element with half axes ω_*xy*_ and ω_*z*_, the autocorrelation function *G*(τ) is defined by 

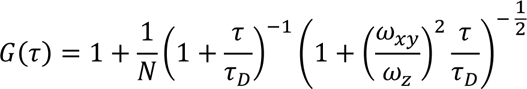

where N is the number of particles, 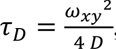, τ*_D_* being the molecular diffusion time of the excited fluorophores moving in a three-dimensional confocal volume through an axial (*z*) to radial (*xy*) dimension, and *D* the diffusion coefficient [cm^2^/s]. Evaluation of the autocorrelated curves was performed with the FCS ACCESS Fit (Carl Zeiss-Evotec) software package using a Marquardt nonlinear least-squares algorithm for a one-component fitting model (Rigler and Elson, 2001). The average hydrodynamic radius R_h_ of the protein was calculated from the obtained translational diffusion coefficient D_trans_ using the Stokes-Einstein relation

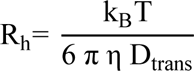

where k_B_ is the Boltzmann constant (1.38 x 10^-23^ J/K), T is the temperature (293 K), and *η* is the viscosity of the solvent (0.001 kg m^-1^ s^-1^). The molecular weight of the protein was estimated by 
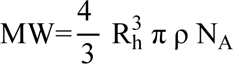
where *N*_A_ is Avogadro’s number = 6.023 × 10^23^ mol^−1^, and *ρ* is the mean density of the molecule. By titrating 40 - 0.2 µM SPOC to 15 nM of Atto-488 labeled peptides pS2 or pS2pS5, the diffusion times significantly increased from 74.4 ± 2.7 µsec (pS2) or 75.0 ± 0.9 µsec (pS2pS5) for peptide alone to 133.0 ± 7.8 µsec (pS2-SPOC) or 127.0 ± 5.0 µsec (pS2pS5-SPOC) for SPOC-bound peptides (Figure S3). The calculated corresponding molar masses of 1.9 ± 0.2 kDa (pS2), 2.0 ± 0.1 kDa (pS2pS5), 17.9 ± 3.6 kDa (SPOC), 20.3 ± 3.8 kDa (pS2-SPOC) and 17.7 ± 2.2 (pS2pS5-SPOC) clearly showed a 1:1 binding stoichiometry (Figure S3C).

### X-ray crystallography

Due to the low sequence identity with published SPOC domain structures of human SHARP and Arabidopsis FPA, PHF3 SPOC structure was solved using the single-wavelength anomalous diffraction (SAD) method. Initial crystals of SPOC at 5 mg/mL were obtained using the sitting-drop vapor diffusion technique and a nanodrop-dispensing robot (Phoenix RE; Rigaku Europe). Crystallization conditions were optimized using microseed matrix screening approach (MMS) (D’Arcy et al., 2014). The best diffracting crystals were grown in conditions B3 from ShotGun HT screen (SG1 HT96 Molecular Dimensions, Suffolk, UK) containing 0.2 M MgCl_2_, 0.1M Bis-Tris pH 5.5, 25% PEG 3350 at 22°C. For co-crystal structures, a 3-fold molar excess of pS2 (Clonestar), pS2pS5 (Clonestar) and pS2pS7 CTD (Eurogentec) peptides was incubated with 5 mg/mL of SPOC. Co-crystals were grown using the sitting-drop vapor diffusion technique. The best diffracting crystals of SPOC:pS2 were obtained using MMS approach in Morpheus screen E9 conditions (Morpheus HT, Molecular Dimensions) containing 0.12 M ethylene glycol mixture, 0.1 M Tris-Bicine buffer pH 8.5, 20% glycerol and 10% PEG 4000; crystals of SPOC:pS2pS7 were obtained in JCSG C4 conditions (JCSG HT96, Molecular Dimensions) containing 0.1 M Hepes pH 7.0, 10% PEG 6000; crystals of SPOC: pS2pS5 were obtained in Morpheus A3 conditions containing 0.03 M MgCl_2_, 0.03 M CaCl_2_, 0.1 M imidazole-MES pH 6.5, 20% glycerol and 10% PEG 4000. The crystals were flash cooled in liquid nitrogen prior to data collection. The selenomethionine data set was collected at the beamline ID29 (ESRF, Grenoble) at 100K at the peak of selenium using a wavelength of 0.979 Å. The data sets of SPOC-CTD peptide complexes were collected at the MASSIF beamline ID30a1 (ESRF, Grenoble) at 100K using a wavelength of 0.966 Å. The data set of SPOC:pS2pS5 was collected at the beamline ID29 (ESRF, Grenoble) using a wavelength 1.07 Å. The data frames were processed using the XDS package (Kabsch, 2010), and converted to mtz format with the program AIMLESS (Winn et al., 2011). The apo-SPOC structure was solved using single anomalous diffraction with the CRANK 2 software suite (Pannu et al., 2011). The structures of SPOC in complex with pS2, pS2pS5 and pS2pS7 CTD peptides were solved using the molecular replacement program PHASER (McCoy et al., 2007) with atomic coordinates of apo-SPOC as a search model. The structures were then refined with REFMAC (Murshudov et al., 1997; Winn et al., 2011) and Phenix Refine (Adams et al., 2010) and rebuilt using Coot (Emsley and Cowtan, 2004). The structures were validated and corrected using PDB_REDO server (Joosten et al., 2014). The figures were produced using the PyMol software. Coordinates were deposited in the protein data bank (accession codes: 6IC8 for PHF3 SPOC:pS2, 6IC9 for PHF3 SPOC:pS2pS7, 6Q2V for PHF3 SPOC, 6Q5Y for PHF3 SPOC:pS2pS5). Data collection and refinement statistics are reported in Table 1. The crystal structure of pS2pS5 CTD peptide bound to PHF3 SPOC showed a different binding mode compared to pS2 and pS2pS7 CTD peptides (Figure S2I-L). Two molecules of SPOC were bound to pS2pS5 peptide, the conformation of the bound peptide was slightly different and three phosphorylated CTD residues (pS5 from the first heptapeptide and pS2pS5 from the second) were forming hydrogen bonds with SPOC residues (Figure S2I-L). SPOC residues K1267 and K1309 from the SPOC_C molecule and R1297 from SPOC_A formed hydrogen bonds with pS5_a_; R1248 from SPOC_C with pS2_b_; and R1248 and R1295 from SPOC_C and K1260 from SPOC_A hydrogen bonded with pS5_b_) (Figure S2L). In sum, both positively charged patches from two SPOC molecules contributed interchangeably to the docking of the three phosphorylated serines (Figure S2L). While pS2 structures indicate that the positive patches on the SPOC surface are geared to accommodate pS2 marks on adjacent repeats, pS2pS5 structure reveals that the binding site can also adjust itself towards binding of phosphomarks within the same repeat.

**Table 1.** X-ray data collection and refinement

|  | **SMet apoSPOC** | **SPOC:2xpS2pS5** | **SPOC:2xpS2** | **SPOC:2xpS2pS7** |
| --- | --- | --- | --- | --- |
| Source ESRF | ID29 |  | MASSIF ID30a1 |  |
| Wavelength (Å) | 0.979 | 1.072 | 0.966 | 0.966 |
| Resolution (Å) | 48.78-2.59<br>(2.71-2.59) | 44.5-2.85<br>(3.0-2.85) | 47.47-1.93<br>(1.97-1.93) | 49.1-1.75<br>(1.78-1.75) |
| Space group | *P*2<sub>1</sub> | *P*3<sub>2</sub> | *P*2<sub>1</sub>2<sub>1</sub>2<sub>1</sub> | *P*2<sub>1</sub>2<sub>1</sub>2<sub>1</sub> |
| Unit cell (Å, °) | 69.75, 69.31,<br>127.22;<br>β=100.09 | 67.83, 67.83,<br>178.11 | 54.07, 68.02,<br>99.17 | 56.22, 68.11,<br>100.87 |
| Molecules (a.u.) | 8 | 4 | 2 | 2 |
| Unique reflections | 32812 (2252) | 21184 (3104) | 27050 (1262) | 39627 (1968) |
| Completeness (%) | 87.8 (50.3) | 98.8 (99.5) | 96.0 (69.0) | 99.2 (91.3) |
| *R*<sub>merge</sub><sup>b</sup> | 0.105 (0.867) | 0.157 (1.432) | 0.076 (1.525) | 0.058 (2.753) |
| *R*<sub>meas</sub><sup>c</sup> | 0.126 (1.178) | 0.183 (1.662) | 0.083 (1.69) | 0.065 (3.143) |
| CC(1/2) | 0.997 (0.520) | 0.993 (0.285) | 0.990 (0.326) | 0.990 (0.230) |
| Multiplicity | 6.0 (3.6) | 3.5 (3.6) | 6.2 (5.1) | 4.4 (4.1) |
| *I*/sig(*I*) | 10.5 (1.1) | 6.6 (0.9) | 11.4 (1.0) | 11.8 (0.5) |
| *B*<sub>Wilson</sub> (Å<sup>2</sup>) | 51.7 | 52.9 | 41.3 | 35.1 |
| *R*<sub>work</sub><sup>e</sup>/*R*<sub>free</sub><sup>f</sup> (%) | 20.6/25.9 | 19.2/22.7 | 18.4/22.6 | 19.5/23.2 |
| r.m.s.d. bonds (Å) | 0.003 | 0.005 | 0.007 | 0.015 |
| r.m.s.d. angles (°) | 0.605 | 1.385 | 0.949 | 1.329 |

<sup>a</sup> Values in parentheses are for the highest resolution shell.
where $\bar{I}_{(hkl)}$ is the mean intensity of multiple $I_{i(hkl)}$ observations of the symmetry-related reflections, *N* is the redundancy
<sup>f</sup> *R*<sub>free</sub> is the cross-validation *R*<sub>factor</sub> computed for the randomly chosen test set of reflections (5 %) which are omitted in the refinement process.

### Immunofluorescence

Cells were grown on glass coverslips, washed with PBS or with PEM buffer (100 mM Pipes, 5 mM EGTA, 2 mM MgCl_2_, pH 6.8) for the neurons and astrocytes and fixed in 4% paraformaldehyde for 10 min. After rinsing with PBS, the cells were permeabilized with a 0.1% Triton X-100 solution in PBS for 8 min, rinsed again and blocked for 1 h RT in blocking buffer (0.1% Tween, 1% BSA in PBS). Coverslips were incubated with the primary antibodies for 1 h RT, washed and subsequently incubated with secondary Alexa-conjugated antibodies for 1 h RT. After washing, coverslips were stained with DAPI and mounted on glass slides in ProLong Gold antifade reagent (Life technologies). Images were acquired using an LSM710 confocal microscope and processed with ImageJ software.

### EU incorporation assay

HEK293T PHF3 WT, KO and ΔSPOC cells were grown for 24 h in 24-well plates for FACS analysis or on coverslips for immunofluorescence. Cells were then incubated with 0.5 µM EU (Molecular Probes) for 1 h. For immunofluorescence, cells were fixed in 2% PFA, washed in 3% BSA in PBS and permeabilized in 0.5% Triton X-100 in PBS. Click-iT^®^ reaction was performed to couple Alexa Fluor 488 Azide (Molecular Probes) to the incorporated EU. Cells were subsequently stained with DAPI and coverslips were mounted on glass slides with ProLong Gold. For FACS analysis, cells were harvested by trypsinization, washed in PBS and fixed overnight in 75% methanol at -20°C. Fixed cells were washed with PBS, blocked in 3% BSA and permeabilized in 0.25% Triton X-100 in PBS. Click-iT reaction was performed to couple Alexa Fluor 488 Azide to the incorporated EU. Cells were subsequently washed twice in 3% BSA in PBS and finally resuspended in PBS. FACS measurements were performed on BD Fortessa machine using Diva software. A population of approximately 10^4^ cells was analyzed for each sample and cell counts in the gated P1 population were measured for three independent experiments using Flowing Software version 2.5.1. Cell counts for each fluorescence intensity were also exported in Microsoft Excel 2010 as frequency distributions of arbitrary fluorescence unit values. Average distributions of three independent experiments were plotted to generate the final FACS data histograms.

### PRO-seq

The protocol was adapted from Kwak et al, 2013. To isolate nuclei, cells were resuspended in cold buffer A (10 mM Tris-Cl pH 8, 300 mM sucrose, 3 mM CaCl_2_, 2 mM MgAc_2_, 0.1% TritonX-100, 0.5 mM DTT), incubated on ice for 5 min and transferred to a dounce homogenizer. After douncing 25 times with the loose pestle, cells were centrifuged at 700 g for 5 min. The pellet was resuspended again in buffer A, centrifuged and resuspended in cold buffer D (10 mM Tris-Cl pH 8, 25% glycerol, 5 mM MgAc_2_, 0.1 mM EDTA, 5 mM DTT), flash frozen with liquid nitrogen and stored at –80°C. For each run-on, 10^7^ HEK293 nuclei was mixed with 10^6^ of Drosophila S2 nuclei (10% for spike-in normalization) in 100 µL buffer D and incubated at 30°C for 3 min with 0.025 mM biotin-11-NTPs and run-on master mix (5 mM Tris-Cl pH 8, 2.5 mM MgCl_2_, 0.5 mM DTT, 150 mM KCl, 0.2 units/µL SUPERase In, 0.5% Sarkosyl). Nascent RNA was isolated using TRIzol LS reagent according to the manufacturer’s instructions, denatured at 65°C for 40 s, hydrolyzed using 0.2 M NaOH on ice for 20 min and neutralized with 1 volume of 1 M Tris-Cl pH 6.8. Buffer was exchanged with DEPC water using BioRad P-30 columns. Fragmented nascent RNA was subsequently enriched using Streptavidin M280 beads by rotating the samples for 20 min in binding buffer (10 mM Tris-Cl 7.4, 300 mM NaCl, 0.1% TritonX-100). Beads were subsequently washed twice with high-salt wash buffer (50 mM Tris-Cl pH 7.4, 2 M NaCl, 0.5% TritonX-100), twice with binding buffer and once with low-salt wash buffer (5 mM Tris-Cl pH 7.4, 0.1% TritonX-100). RNA was isolated from the beads using TRIzol reagent in two consecutive rounds and pooled together for ethanol precipitation. The RNA pellet was redissolved in DEPC H_2_O with 10 pmol of reverse 3’ RNA adaptor starting with a 5’ random octamer sequence (5’-5Phospho rNrNrNrNrNrNrNrNrGrArUrCrGrUrCrGrGrArCrUrGrUrArGrArArCrUrCrUrGrArArC-/inverted dT/ -3’) and subjected to ligation using T4 RNA ligase I (NEB) at 16°C ON. RNA was isolated using Streptavidin M280 beads as previously described and 5’ ends were repaired using Cap-Clip Acid Pyrophosphatase (Cellscript) and Polynucleotide Kinase (PNK, NEB) according to manufacturers’ instructions. RNA was purified again with TRIzol and ethanol precipitation as before and subjected to 5’ RNA adaptor ligation as for the 3’ adaptor ligation (the 5’ adaptor contained a 3’ random tetramer sequence 5’-rCrCrUrUrGrGrCrArCrCrCrGrArGrArArUrUrCrCrArNrNrNrN -3’). RNA was enriched by a third round of binding to Streptavidin M280 beads and TRIzol isolation. RNA was retro-transcribed using RP1 Illumina primer and SuperScript III (Invitrogen) to generate cDNA libraries. Libraries were then amplified using KAPA HiFi Real-Time PCR Library Amplification Kit (Peqlab) and Illumina primers containing standard TruSeq barcodes. Amplified libraries were subjected to electrophoresis on 2.5% low melting agarose gel and amplicons from 150 to 300 bp were excised, purified from the gel using NucleoSpin Gel and PCR Clean-up kit (Macherey-Nagel) and sequenced on Illumina HiSeq 2500 platform (VBCF NGS facility).

### Transcription elongation inhibition with DRB and release

PHF3 WT, KO and ΔSPOC HEK293T cells were grown in 15 cm dishes and incubated with 100 µM DRB (Sigma) or DMSO for 3.5 h. Cells for time point 0 were harvested immediately. Alternatively, cells were washed twice with PBS and allowed to recover in normal medium for 10, 25 or 40 min before harvesting. Cells were harvested and immediately processed for nuclei isolation as described in the PRO-seq section.

### RNA isolation, RT-qPCR and RNA-seq library preparation

RNA was isolated from harvested cells using TRI reagent (Sigma) according to the manufacturer’s instructions. cDNA was obtained by reverse transcription of 1 µg of RNA using random hexamer primers (Invitrogen) and ProtoScript II Reverse Transcriptase (NEB) according to manufacturers’ instructions. RT-qPCR was performed on a BioRad CFX384 Touch qPCR cycler using iTaq Universal Sybr Green Supermix (BioRad). RT-qPCR data were analyzed by normalising the expression of the genes of interest by GAPDH housekeeping gene expression; gene expression was calculated after assessing primers efficiency. For RNA-seq, 8x10^6^ HEK293 cells were mixed with 2x10^6^ Drosophila S2 cells for spike-in normalization. Total isolated RNA was first treated with recombinant DNaseI (Roche), cleaned up using peqGOLD PhaseTrap A tubes (Peqlab), and rRNA-depleted using the Ribo-Zero kit (Illumina). RNA-seq libraries were prepared using the NEBNext Ultra II directional RNA library preparation kit (NEB) according to the manufacturer’s instructions. Sequencing was performed on Illumina HiSeq 2500 (VBCF NGS facility).

### Chromatin immunoprecipitation

Cells were harvested, counted, resuspended in 50 mL PBS/10^8^ cells and fixed for 10 minutes by adding formaldehyde to a final concentration of 1%. Formaldehyde was quenched by adding glycine pH 3 to a final concentration of 0.6 M for 15 minutes. Cells were centrifuged and washed twice in cold PBS. To isolate nuclei, 10^8^ fixed cells were resuspended in 5 mL cold lysis buffer 1 (50 mM Hepes/KOH pH 7.5, 140 mM NaCl, 1 mM EDTA, 10% glycerol, 0.5% Nonidet P-40, 0.25% Triton X-100, 1x protease inhibitor; for Pol II ChIP 2 mM Na_3_VO_4_ and 2 mM NaF), rotated for 10 min at 4°C and centrifuged. Nuclei were resuspended in 5 mL cold lysis buffer 2 (10 mM Tris-Cl pH 8, 200 mM NaCl, 1 mM EDTA, min at room temperature and centrifuged. The pellet was resuspended in 3 mL lysis buffer 3 (10 mM Tris-Cl pH 8, 100 mM NaCl, 1 mM EDTA, 0.5 mM EGTA, 0.1% Na-deoxycholate, 0.5% N-lauroylsarcosine, 1x protease inhibitors; for Pol II ChIP 2 mM Na_3_VO_4_ and 2 mM NaF). Chromatin was sheared to an average size of 200-600 bp using the Bioruptor Pico (Diagenode) for 20 cycles, 30 sec on/30 sec off. Triton X-100 was added to a final concentration of 1%. 5-10% of chromatin was kept as an input, to the remainder antibody (Pol II pS5 3E8; Pol II pS2 3E10; Pol II pS7 4E12; total Pol II clone F-12 Santa Cruz sc-55492; TCEA1 Abcam ab185947; H3K27me3 Millipore 07-449) or antiserum (GFP Abcam ab290) was added and rotated ON at 4°C. Antibody and cell amounts are indicated in the key ressources table. For TFIIS, Pol II F-12, and H3K27me3 ChIP, chromatin was mixed with 2.5% of mouse chromatin as a spike-in before adding the antibody. Protein G or protein A Dynabeads were washed three times in cold block solution (0.5% BSA in PBS), antibody-bound chromatin was added to the beads and rotated 4-6 hours at 4°C. Beads were washed 5 times (8 times for Pol II pS5 ChIP) in RIPA washing buffer (50 mM Hepes/KOH pH 7.5, 500 mM LiCl, 1 mM EDTA, 1% NP-40, 0.7% Na-deoxycholate) and once in 50 mM NaCl in TE. Crosslinked protein-DNA complexes were eluted in 200 µL elution buffer (50 mM Tris-Cl pH 8, 10 mM EDTA, 1% SDS) for 15 min at 65°C. Crosslinks were reversed at 65°C ON. RNA was degraded by adding 0.2 mg/mL RNase A for 2 h at 37°C, proteins were digested by adding 0.2 mg/mL proteinase K and 5.25 mM CaCl_2_ for 30 min at 55°C. DNA was purified by phenol-chloroform extraction, ethanol-precipitated and resuspended in 50 µL nuclease-free water. Next generation sequencing libraries were prepared using the NEBNext Ultra II DNA library Prep Kit for Illumina and NEBNext Multiplex Oligos Primer Set 1-3 (New England Biolabs) according to the manufacturer’s instructions. Next generation sequencing was performed on Illumina HiSeq 2500, NextSeq 550 or NovaSeq 6000 (VBCF NGS facility).

### Pol II phosphorylation, elongation complex (EC) preparation and sucrose gradient ultracentrifugation

Pol II was phosphorylated with DYRK1A kinase (generously provided by Matthias Geyer) in kinase buffer (50 mM Hepes pH 7.5, 34 mM KCl, 7 mM MgCl_2_, 5 mM β-glycerophosphate, 2.5 mM DTT) with 1 mM ATP for 1 h at 30°C. A nucleic acid scaffold for transcribing Pol II was assembled by mixing equimolar amounts of DNA (EC-phf3-template) and RNA (RNA50) in a BioRad T100 Thermal Cycler heated to 95°C and cooled in 0.1°C/s increments until 4°C was reached. For sucrose gradient ultracentrifugation, the Pol II-EC was assembled by incubating 60 pmol Pol II with a 2-fold molar excess of DNA/RNA for 10 min on ice, followed by 10 min at 30°C, and another 10 min at 30°C after adding a 4-fold molar excess of non-template DNA (EC-phf3-nontemplate) to generate a transcription bubble. A 4-fold molar excess of PHF3 or TFIIS^M^/TFIIS^M^+TFIIF was incubated with Pol II-EC for 20 min at 25°C, followed by addition of the 4-fold molar excess of the competitor (TFIIS^M^/TFIIS^M^+TFIIF or PHF3 respectively) for 20 min at 25°C. 10-30% sucrose gradients were prepared using a gradient mixer (Gradient Master 108; BioComp Instruments). Pol II complexes were applied on top of the gradient followed by ultracentrifugation at 32 000 rpm in a SW60 swinging bucket rotor (Beckman Coulter) for 16 h at 4°C. 80 µl fractions were collected carefully from top to the bottom of the tube and analyzed by Western blotting.

### *In vitro* transcription elongation assay

Pol II phosphorylation and transcription bubble assembly were performed as above, using arrest sequences comprising a region for EC assembly and a region containing a previously characterized Pol II arrest site shown to be responsive to TFIIS (Arrest-template/5’-FAM-Arrest-RNA and Arrest-nontemplate) (Hawryluk et al., 2004). 0.12 µM Pol II was used per reaction. 0.1 µM TFIIS and different concentrations of PHF3 or PHF3 ΔTLD were added to Pol II-EC and incubated for 5 min at 30°C. Transcription was initiated by adding 100 µM of NTPs in a transcription buffer (20 mM Hepes pH 7.5, 75 mM NaCl, 3 mM MgCl_2_ and 4% glycerol) and incubating at 37°C for 10 min. Final ATP concentration was 200 µM due to the leftover from the kinase reaction. Reaction were stopped by adding urea loading buffer (4M urea in TBE) and EDTA (12.5 mM) and boiling at 95°C for 5 min. After chilling on ice, samples were incubated with 0.1 mg/mL proteinase K at 37°C for 20 min, boiled at 95°C for 5 min and chilled on ice before loading on a 20% denaturing acrylamide gel. Gels were run at 300 V for 1.5 h and scanned using Typhoon (GE Healthcare).

### SLAM-seq

SLAM-seq was performed as described (Herzog et al., 2017). HEK293T cells seeded into 6 cm dishes the day before the experiment were incubated in standard culture medium containing 100 µM 4-thiouridine (s^4^U; ChemGenes) for 12 h with media exchanges every 3 hrs. Subsequently, cells were washed twice with PBS and incubated with 10 mM uridine-containing medium. Cells were harvested using TRI reagent (Sigma) at timepoints 0 h, 6 h and 12 h after removal of s^4^U. RNA was isolated according to the manufacturer’s instructions including 0.1 mM DTT during isopropanol precipitation and resuspended in 1 mM DTT. Isolated RNA was treated with 10 mM iodacetamide to alkylate s^4^U and subjected to ethanol precipitation. Alkylated RNA was resuspended in water and treated with TURBO DNA-free Kit (Invitrogen) according to the manufacturer’s instructions. For global SLAM-seq, libraries were prepared using QuantSeq 3’ mRNA-Seq Library Prep Kit FWD for Illumina (Lexogen) according to the manufacturer’s instructions. For targeted analysis of INA, RNA was reverse transcribed using oligo(dT) primer. Subsequently, RNA was removed using RNAse H (NEB) according to the manufacturer’s instructions and cDNA was amplified using INA-specific forward primer (5’-CACGACGCTCTTCCGATCTNNNNNNTCTGTCCAGCAGTCACTTCG-3’) and oligo(dT) primer (5’-GTTCAGACGTGTGCTCTTCCGATCT-(T)n-V-3’) for 23 cycles. Library amplification was performed using Illumina FWD (5’-AATGATACGGCGACCACCGAGATCTACACTCTTTCCCTACACGACGCTCTTCCGATCT-3’) and REV (5’-CAAGCAGAAGACGGCATACGAGATNNNNNNGTGACTGGAGTTCAGACGTGTGCTCTTCCGATCT-3’) index primers. Next generation sequencing was performed at the VBCF NGS facility using Illumina NextSeq 550.

### Genomic region definition

To reliably quantify gene activity, transcription start sites (TSS) and gene body regions were precisely defined. TSS for HEK293 cells were extracted from the FANTOM5 data set (24670764), and transferred from hg19 to the hg38 version of the human reference genome using liftOver (16381938). Regions which mapped to multiple locations were disregarded. Each Fantom TSS was extended into a promoter region of +/-250 bp from the putative end of the TSS region. Gene body regions were defined in the following way. Firstly, gene models were downloaded from the Ensembl database (27899575) on 02.10.2015. Each promoter region was assigned to the nearest transcript on the corresponding strand from the Ensembl annotation; promoters that were >2 kb away from a transcript were removed from the data set. Gene body region was defined as a region from the promoter end (+250 bp from the TSS) to the most commonly annotated transcript end for the corresponding gene (a transcript end which is supported by highest number of annotated isoforms). If multiple transcript ends had the same support, the longest isoform was chosen as the representative. If the corresponding transcript overlapped with multiple defined TSS regions, a representative promoter was chosen for each gene by selecting the TSS with the highest average PRO-seq signal.

### Analysis of PRO-seq data

Prior to quantification of PRO-seq data, genomic regions containing genes were split into promoter regions and gene bodies. PRO-seq data were mapped to the hg38 version of the human reference genome using the STAR-2.4.0 (23104886) aligner with the following parameters: --outFilterMultimapNmax 10 --outFilterMismatchNoverLmax 0.2 --sjdbScore 2. PRO-seq signal in the promoter area was quantified by counting the number of read 5’ ends overlapping with the defined promoter regions, while the PRO-seq signal within the gene body was quantified by counting the number of read 5’ ends overlapping the gene body. PRO-seq counts for each region and each sample were normalized to Drosophila S2 spike-in by multiplying the corresponding counts with the ratio between the total number of human and Drosophila reads. Differential analysis was performed using DESeq2 (Love et al., 2014). Spike-in calculated normalization factors were normalized to have geometric mean of 1, prior to the differential analysis. To avoid the possibility of quantifying transcriptional read-through from highly expressed genes, all genes which had wild type PRO-seq RPKM > 8 in a 250 bp region 250 bp upstream of the promoter region were filtered out of the analysis. Additionally, all non-expressed genes (genes with promoter RPKM < 2 in all samples) were also filtered from the analysis. PRO-seq data was deposited under the accession number E-MTAB-7501.

### Validation of PRO-seq data

The quality of PRO-seq data was evaluated using the WT sample, which showed that the most highly expressed genes are microRNAs, histones, Pol II and Jun, as already reported. We compared the expression values in the TSS regions between FANTOM5 data based on Cap Analysis of Gene Expression (CAGE) of mRNAs and our PRO-seq data containing nascent RNAs (Figure S7). We obtained a high Spearman correlation between our data and the CAGE data (0.57) and we observed a spread in the PRO-seq data where CAGE data have low signal, indicating that PRO-seq captures the signal in a much lower dynamic range. In addition, we compared the PRO-seq data with the NET-seq from HEK293 cells for the TSS regions (Mayer et al., 2015). NET-seq data bigWig files were downloaded from the GEO database (GSE61332), and transferred from the hg19 to hg38 genome versions using CrossMap (Zhao et al., 2014). The NET-seq signals were summarized using the Bioconductor package genomation and compared to the average normalized PRO-seq signal using the Spearman correlation coefficient. The correlation was visualized using ComplexHeatmap (Gu et al., 2016). The Spearman correlation between PRO-seq and NET-seq samples was >0.85, which confirms that the data are of high quality. The heatmap shows the correlations between PRO-seq data for PHF3 WT, KO, ΔSPOC, and NET-seq data (Figure S7B).

### Analysis of ChIP-seq data

PHF3, Pol II pS2, pS5, and pS7 ChIP-seq data were mapped to the hg38 version of the human genome using Bowtie2 (Langmead and Salzberg, 2012) with the following parameters: bowtie2 -k 1. ChIP-seq data was deposited under the accession number E-MTAB-8783. PHF3 occupancy positively correlates with Pol II occupancy (TSS Spearman correlation with pS2 is 0.38, with pS5 0.31, with pS7 0.34; gene body correlation with pS2 is 0.74, with pS5 0.68, with pS7 0.72; Figure S4A).

TFIIS, Pol II (F-12), and H3K27me3 ChIP-seq samples were normalized using mouse chromatin spike-in and mapped separately to the human (hg38) and mouse (mm10) genomes, using Bowtie2, as implemented in PigX pipeline (Wurmus et al., 2018). The scaling factor was obtained by dividing the total number of uniquely mapped reads to the human genome, with the number of uniquely mapping reads to the mouse genome. The genomics tracks were then constructed by extending the reads to 200 bp into 3’ direction, calculating the coverage vector, and scaling using the aforementioned scaling factor. TFIIS, Pol II (F-12), and H3K27me3 ChIP-seq data was deposited under the accession number E-MTAB-8789.

### Construction of genomic tracks

Genomic tracks were constructed by merging all replicates of the corresponding biological conditions (WT, KO, ΔSPOC) and experiments (RNA-seq, PRO-seq, ChIP-seq, SLAM-seq). Merged tracks were then normalized to the total number of reads. For ChIP-seq, the reads were firstly extended to 200 bp towards the 3’ end, the coverage was calculated, and normalized to the total number of reads. The tracks were additionally normalized by taking the log2 ((ChIP +1) / (Input+1)). Negative values were censored to zero.

### Signal profile construction

Signal profiles were constructed by averaging the signal from genomic tracks over different functional regions into 100 bins of equal size. The extreme values in the profiles were avoided by applying the trimmed mean function, with the trim parameter set to 0.3. To compare ChIP-seq profiles of different antibodies (different proteins), the profiles were normalized prior to averaging to the within region signal range by dividing the signal by min – max.

### Analysis of HEK293T RNA-seq data

RNA-seq reads were mapped to a genome comprising of human reference genome hg38 version and Drosophila melanogaster version dm6. STAR-2.5.3 was used with the default parameters and gencode v28 gtf annotation as gtf file. RNA-seq data were quantified using STAR quantMode. Differential expression was analyzed using DESeq2 and the gene counts were normalized to the total Drosophila spike-in counts; genes with an adjusted p-value < 0.05 were designated as differentially expressed. HEK293T RNA-seq data was deposited under the accession number E-MTAB-7498.

### Analysis of mES RNA-seq data

Mouse embryonic stem cell RNA-seq data were mapped to the mm9 version of the mouse reference genome using STAR-2.4.0. STAR index was constructed with gene annotation downloaded from the Ensembl database on 20.05.2015. The expression was quantified and the differential expression was analyzed as described previously. mESC RNA-seq data was deposited under the accession number E-MTAB-7526.

Raw sequenced reads were processed with the SLAMdunk pipeline as previously described (Herzog et al., 2017). Genes which had detectable conversion rates in at least three biological replicates in all conditions were kept for subsequent analysis. Furthermore, genes which had a non-monotonic decrease of the median conversion rates were filtered out. To estimate the half-lives, a robust linear model was fit on the linearized form of the exponential decay equation using the RLM function from the MASS R package. SLAM-seq data was deposited under the accession numbers E-MTAB-7898 and E-MTAB-7899.

### Sequencing data integration

The complete data integration and data analysis were done in R using Bioconductor (Huber et al., 2015), and the following libraries: GenomicAlignments (Lawrence et al., 2013), data.table (Matt Dowle and Arun Srinivasan, 2017), data.table: Extension of ‘data.fram’ (R package version 1.10.4-3.), biomaRt (Durinck et al., 2009), GenomicRanges, rtracklayer (Lawrence et al., 2009), SummarizedExperiment (10.18129/B9.bioc.SummarizedExperiment), genomation (Akalin et al., 2015), and ggplot2 (10.1007/978-0-387-98141-3).

### Differentiation of mESCs into neural stem cells, neurons and astrocytes

mESCs differentiation into neural stem cells (NSCs), and later in neurons and astrocytes, was adapted from a previously described protocol (Pollard et al., 2006). Briefly, 10^4^ mESCs/cm^2^ were seeded on gelatin-coated 10 cm dishes and cultured for 7 days in N2B27 medium. After 7 days, 2-5x10^6^ cells were transferred to non-gelatinized T75 flasks in NS-N2B27 medium (N2B27 medium supplemented with 10 ng/mL EGF and 10 ng/mL FGF2) and grown for 2-4 days to form aggregates in suspension. The cell aggregates were then collected by centrifugation (700 rpm for 30 s) and transferred to fresh gelatin-coated T75 flasks and grown in NS-N2B27 medium. After 3 to 7 days, cells displayed NSCs morphology. For neuronal differentiation, NSCs were seeded in NS-N2B27 medium at a density of 25000 cells/cm^2^ on laminin-coated glass coverslips in 24-well plates for immunofluorescence and 6-well plates for RNA isolation. The day after, the medium was replaced with N2B27 medium supplemented with only 5 ng/mL FGF2. Cells were then fixed or harvested at the indicated time points. For astrocyte differentiation, NSCs were seeded in N2B27 medium supplemented with 1% FBS at a density of 50000 cells/cm^2^ on gelatin coated glass coverslips in 24-well plates. Cells were fixed after 5 days and samples were processed for immunofluorescence.

### Exit from naïve pluripotency assay of mESCs

mESCs were cultured in N2B27 medium (DMEM F12 + Neurobasal medium supplemented with 1% L-glutamine, 1% penicillin/streptomycin, 1% NEAA, B27 supplement, N2 supplement, 2-mercaptoethanol) supplemented with 2i/LIF as previously described (Leeb et al., 2015) for at least two passages. 10^4^ cells were seeded in 6-well plates and grown for 36 h in N2B27medium + 2i in presence or absence of LIF. Subsequently, cells that were grown without LIF were incubated in the absence of 2i to allow differentiation for 8 h and 24 h. After harvesting the cells, RNA was isolated to determine gene expression by RT-qPCR.

## QUANTIFICATION AND STATISTICAL ANALYSIS

Error bars represent standard deviation estimated from three to four independent experiments. For Western blot band intensity, EU fluorescence intensity and RT-qPCR analyses statistical significance was calculated using one-tailed Student’s t-test. P-values smaller than 5% were considered statistically significant and indicated with an asterisk (‘*’ for p<0.05; ‘**’ for p<0.01; ‘***’ for p<0.001; ‘****’ for p<0.0001). ChIP-seq, RNA-seq and PRO-seq were performed in triplicates. SLAM-seq was performed in six replicates. Statistical analysis on all sequencing data was performed using DESeq2. Statistical significance of the differential expression/abundance was determined by two-tailed Wald test, after appropriate normalization for each type of data.

## DATA AND SOFTWARE AVAILABILITY

Atomic coordinates were deposited in the Protein Data Bank with accession codes: 6IC8 for PHF3 SPOC:pS2, 6IC9 for PHF3 SPOC:pS2pS7, 6Q2V for PHF3 SPOC, 6Q5Y for PHF3 SPOC:pS2pS5. All sequencing data from this study were deposited in ArrayExpress and are accessible through the accession numbers E-MTAB-7498 (RNA-seq HEK293T), E-MTAB-8783 (PHF3, Pol II pS2, pS5, pS7 ChIP-seq), E-MTAB-8789 (Pol II F-12, TFIIS, H3K27me3), E-MTAB-7501 (PRO-seq), E-MTAB-7898 and E-MTAB-7899 (SLAM-seq), E-MTAB-7526 (RNA-seq mESC).

## SUPPLEMENTAL FIGURES

**Figure S1.**
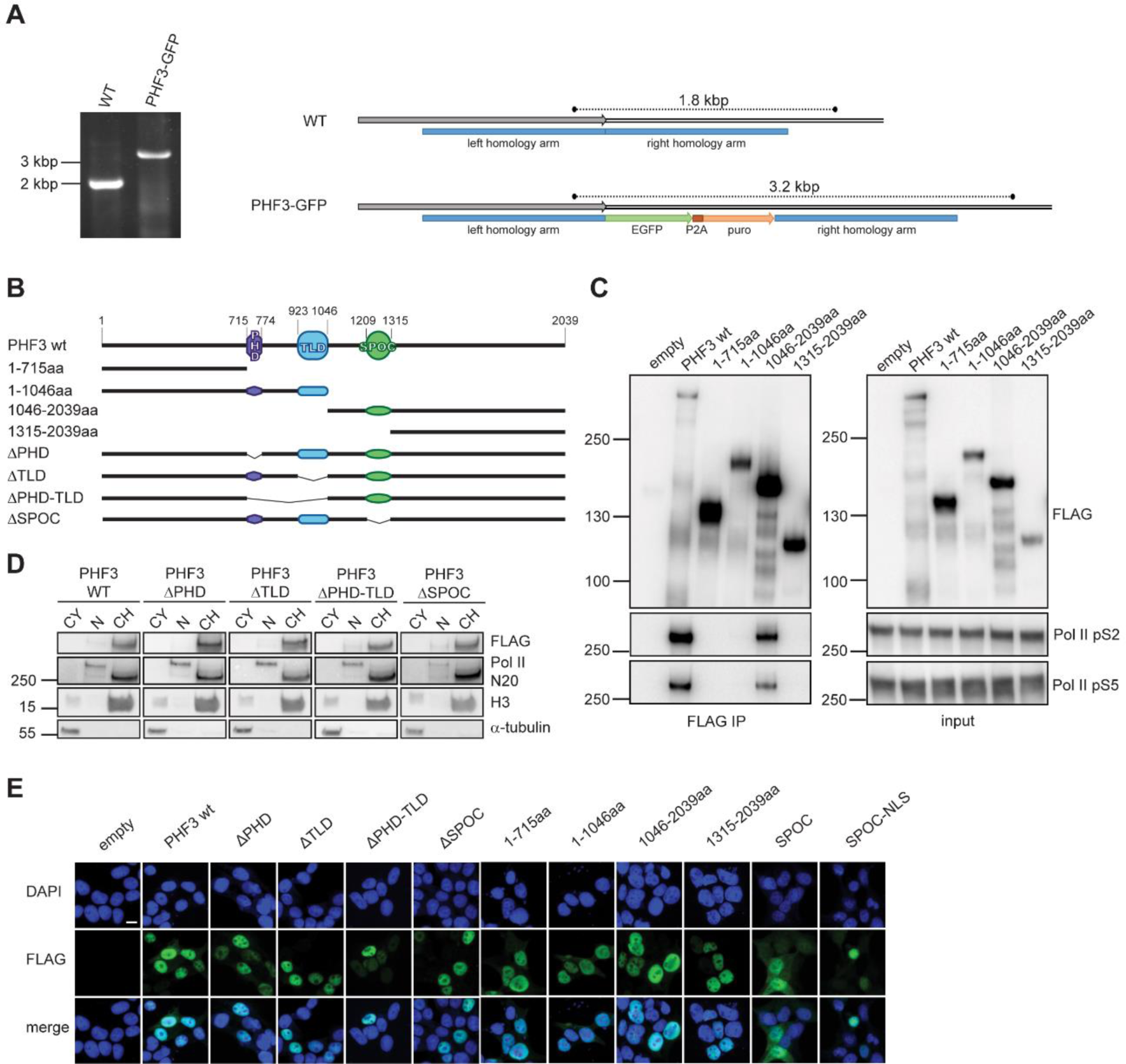
Generation of the PHF3-GFP cell line and immunoprecipitation and localization of PHF3 truncation mutants. (A) CRISPR/Cas9 strategy and PCR validation of endogenous tagging of PHF3 with GFP at the C-terminus. (B) A scheme of PHF3 truncation constructs used in (C-E). (C) Anti-FLAG immunoprecipitation of PHF3 truncation mutants shows loss of interaction with Pol II in the absence of the SPOC domain. (D) Chromatin localization of FLAG-PHF3 truncation mutants revealed by biochemical fractionation and Western blotting. Alpha-tubulin, Pol II and H3 were used as markers of cytoplasmic (CY), nucleoplasmic (N) and chromatin (CH) fractions, respectively. (E) Nuclear localization of FLAG-PHF3 truncation mutants revealed by immunofluorescence staining with anti-FLAG and nuclei counterstaining with DAPI. Scale bar=10 μm.

**Figure S2.**
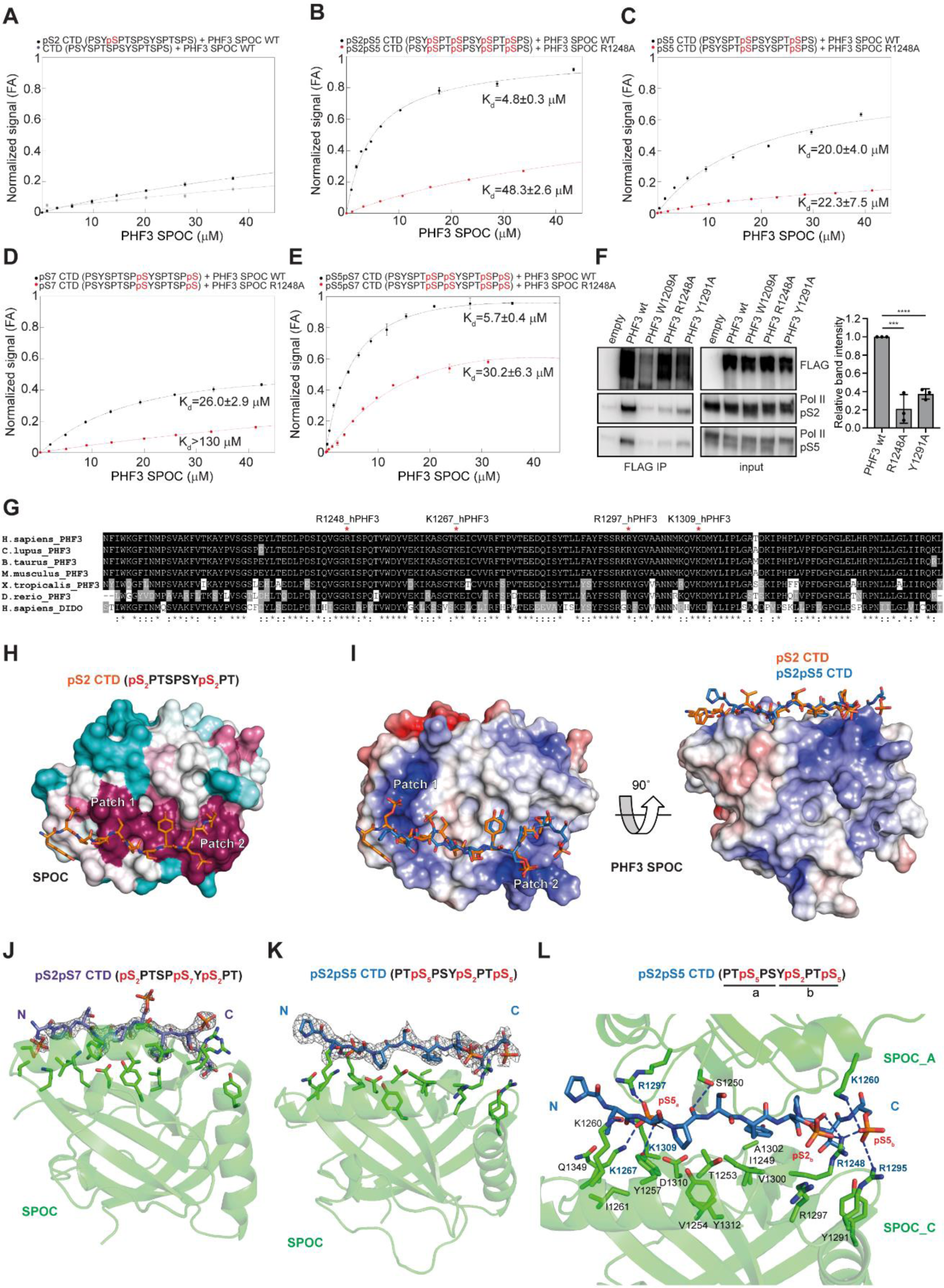
Analysis of interaction between PHF3 SPOC and CTD peptides. (A-E) Fluorescence anisotropy (FA) measurement of the binding of (A) pS2 on one CTD repeat and non-phosphorylated CTD, (B) pS2pS5, (C) pS5, (D) pS7 and (E) pS5pS7 CTD peptides to PHF3 SPOC WT (black) or R1248A mutant (red). Normalized fluorescence anisotropy is plotted as a function of protein concentration (n=3). The data were normalized for visualization purposes and the experimental isotherms were fitted to one site saturation with non-specific binding model. (F) Mutation of conserved residues in PHF3 SPOC shows loss of interaction with Pol II. W1209 is important for structural integrity; R1248 and Y1291 were shown to mediate SHARP SPOC binding to SMART and NCoR peptides (Ariyoshi and Schwabe, 2003). The bar graph shows quantification of Pol II in the eluates, normalized to the immunoprecipitated PHF3 bait (mean±sd; n=3). (G) Multiple sequence alignment of PHF3 SPOC from *Homo sapiens*, *Canis lupus*, *Bos taurus*, *Mus musculus*, *Xenopus tropicalis* and *Danio rerio*, and DIDO SPOC from *Homo sapiens*. Four basic residues shown to bind pS2 CTD in PHF3 SPOC are indicated with a red asterisk. (H) Evolutionary conservation of PHF3 SPOC residues projected onto the pS2 co-structure using the ConSurf server. Residues are colored by their conservation grades with maroon showing the highest and turquoise the lowest degree of conservation. Two positively charged patches (Patch 1 and 2) are indicated. (I) Overlay of PHF3 SPOC structures with pS2 and pS2pS5 shows a difference in conformation of the N-terminal CTD residues. The color coded electrostatic surface potential of SPOC was drawn using the Adaptive Poisson-Boltzmann Solver package within PyMol. The electrostatic potential ranges from −5 (red) to +5 (blue) kT/e. (J) 2 *F_o_* − *F_c_* electron density map of pS2pS7 peptide contoured at the 1.5σ level. (K) 2 *F_o_* − *F_c_* electron density map of pS2pS5 peptide contoured at the 1.5σ level. (L) Hydrogen bonding interactions between pS2pS5 and two PHF3 SPOC molecules. SPOC residue labels from two positively charged patches are colored blue.

**Figure S3.**
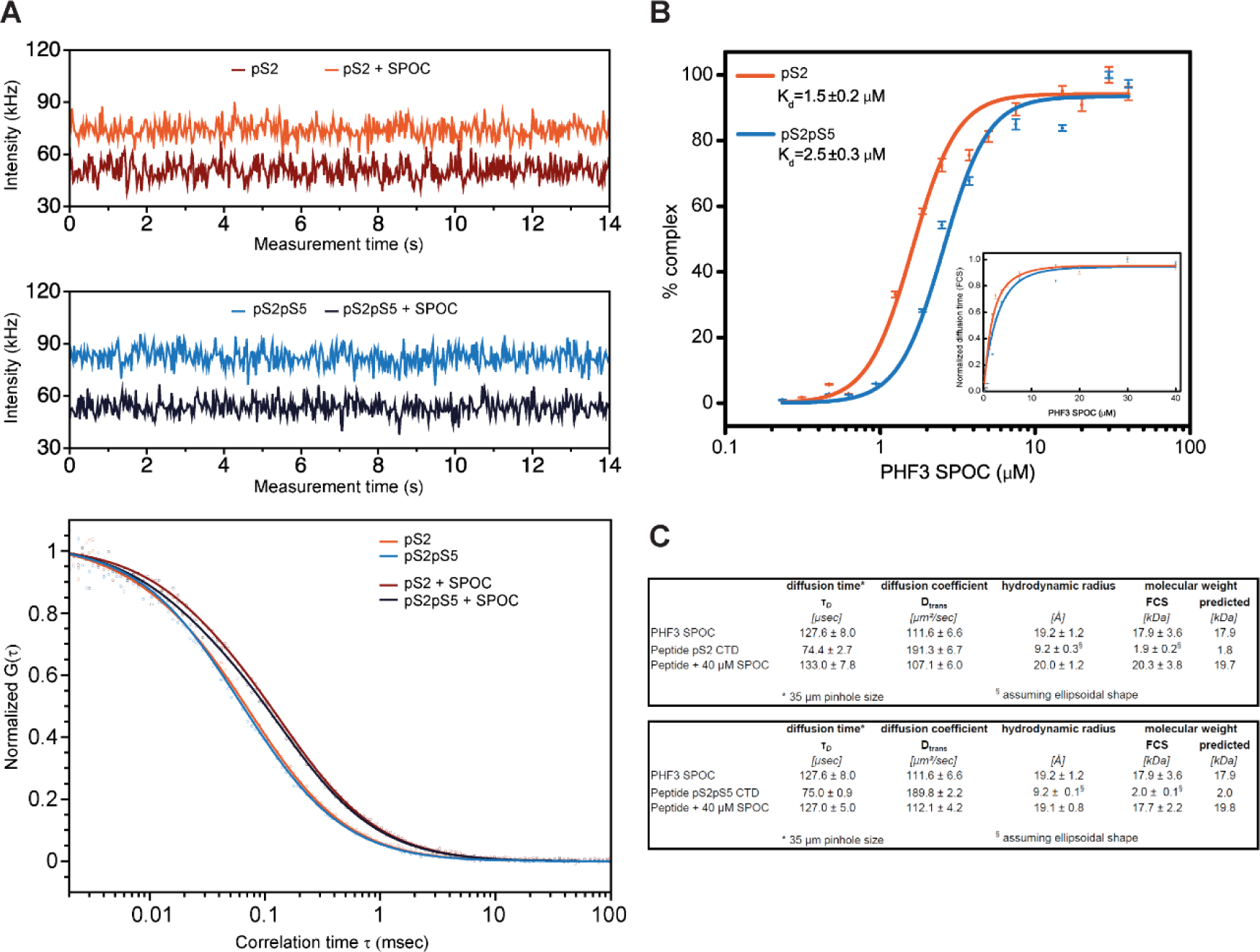
Analysis of binding stoichiometry between PHF3 SPOC and pS2 or pS2pS5 CTD peptides by fluorescence correlation spectroscopy (FCS) (A) Fluorescence intensity fluctuations of all the molecules through a Gaussian confocal volume of 0.65 femtolitres are recorded over 15 seconds. Normalized autocorrelation function G(τ) describes the observed fluctuations of the fluorescence intensity from the mean intensity at any time compared to fluctuations at delayed time. Diffusion parameters (τ*_D_* and D_trans_) are obtained from the normalized autocorrelated dataset using a Marquardt nonlinear least-square fitting routine. (B) Percentage of SPOC-CTD peptide complex formation calculated from the change in translational diffusion behavior. K_d_ values of pS2 and pS2pS5 CTD peptides binding to SPOC are obtained from normalized translational diffusion times (inset). (C) Biophysical parameters of SPOC, pS2 and pS2pS5 peptides and their complexes obtained by FCS (T=20°C).

**Figure S4.**
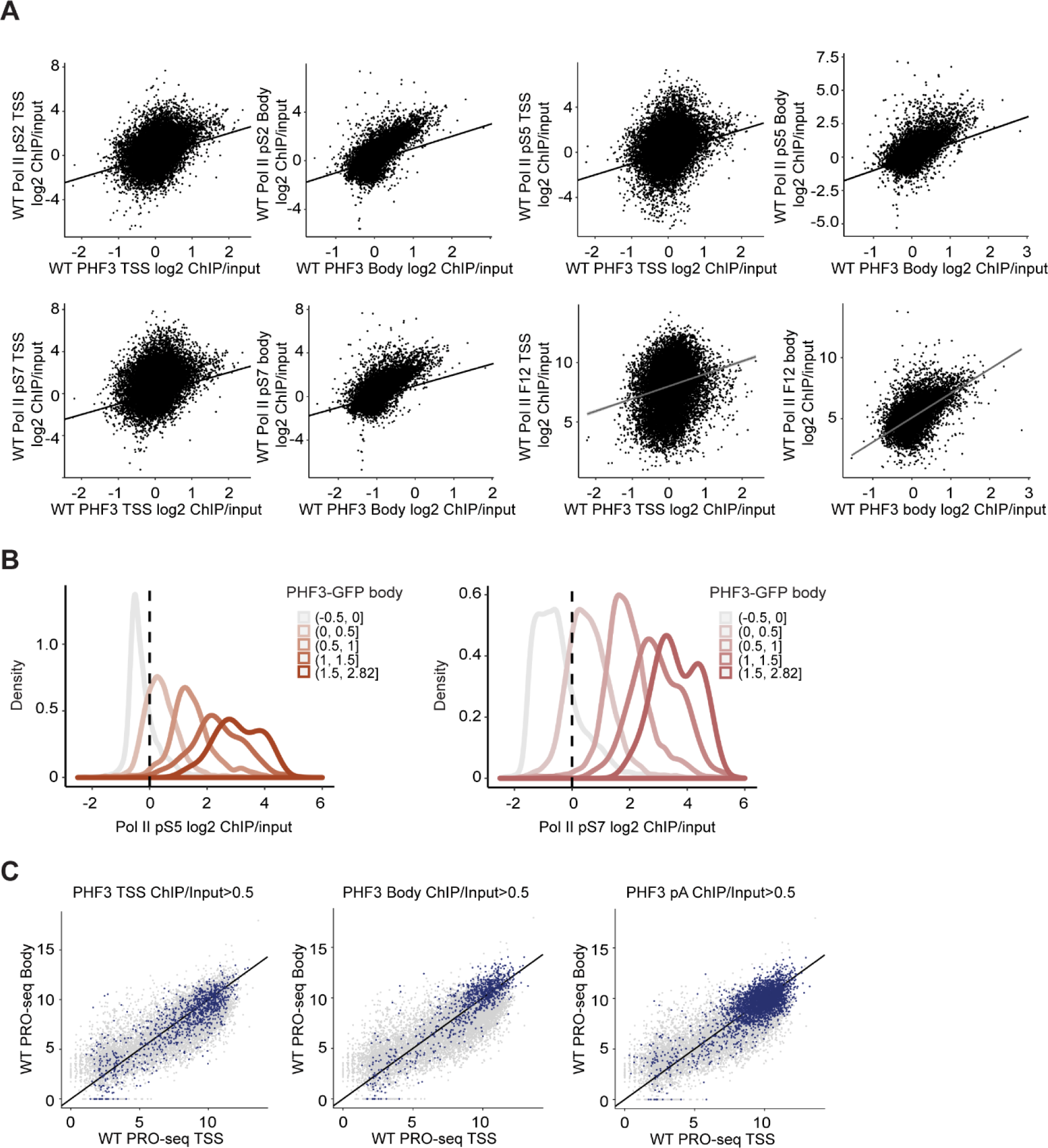
PHF3 genomic occupancy coincides with Pol II and increases with transcription levels. (A) Pol II pS2, pS5, pS7 and F-12 occupancy relative to PHF3 occupancy on TSS and gene body. (B) Pol II pS5 and pS7 log2 ChIP/input in PHF3 WT colored by PHF3 gene body occupancy, with light color showing the lowest and dark color the highest PHF3 occupancy category. (C) Scatter plots showing PRO-seq nascent transcription levels at gene body relative to TSS in WT cells. Blue dots indicate PHF3-bound genes at TSS (left), gene body (middle) or pA (right).

**Figure S5.**
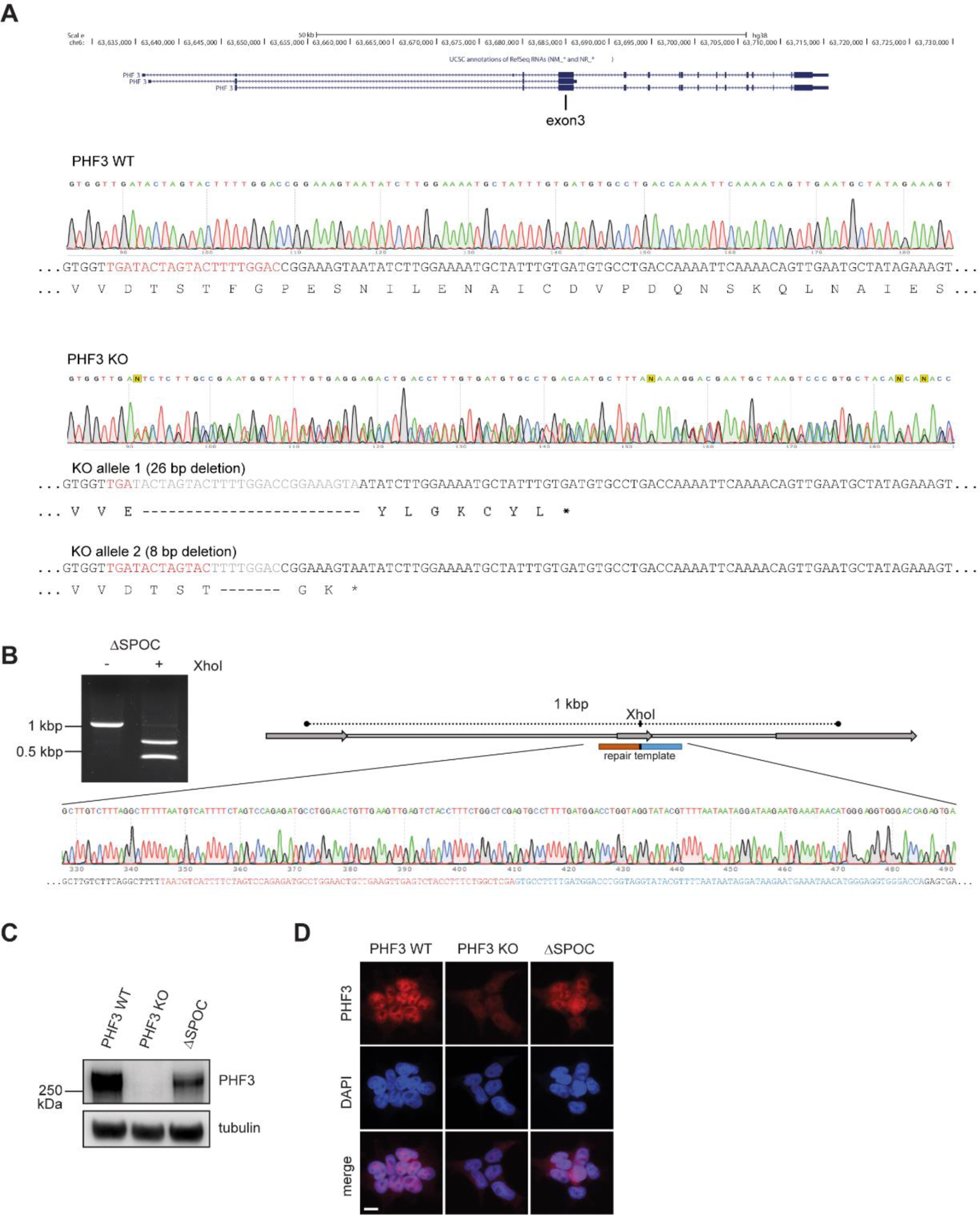
Generation of HEK293T PHF3 knock-out (KO) and ΔSPOC cells by CRISPR/Cas9. (A) Sequencing snapshots and sequence reconstruction of human PHF3 KO in HEK293T cells generated by a frameshift-induced stop codon in exon 3. gRNA target region is shown in red, deleted regions are in grey. (B) Strategy and validation of endogenous deletion of the SPOC domain from PHF3 (ΔSPOC) by PCR and sequencing. (C) Western blot and (D) immunofluorescence analysis of PHF3 KO and ΔSPOC in HEK293T cells.

**Figure S6.**
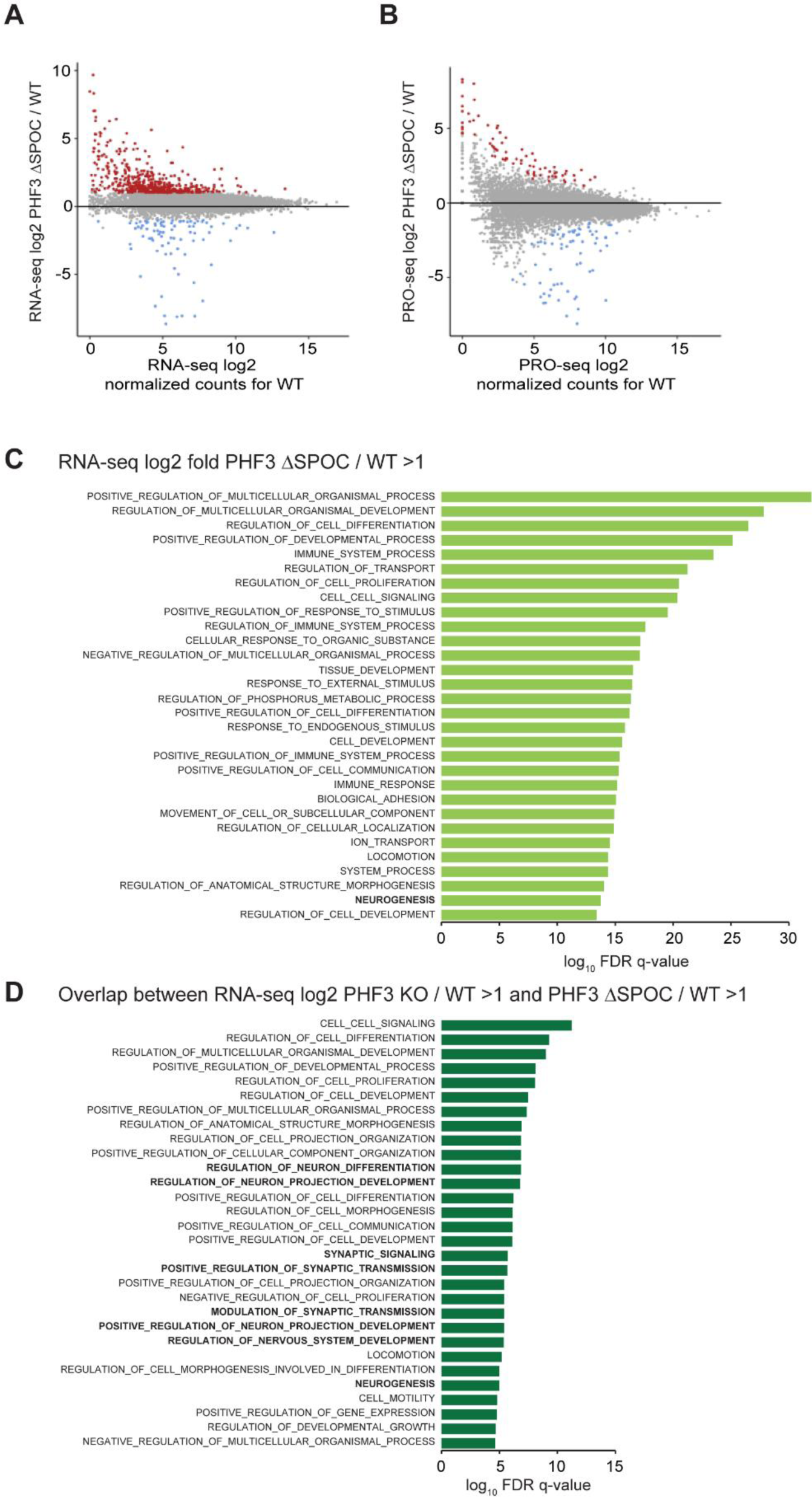
RNA-seq and PRO-seq analysis. (A) RNA-seq analysis shows upregulation of 638 genes (red dots, fold-change>2, p<0.05) and downregulation of 74 genes (blue dots, fold-change>2, p<0.05) in PHF3 ΔSPOC compared to WT. Drosophila S2 cells were used for spike-in normalization. (B) PRO-seq analysis shows upregulation of 78 genes (red dots, fold-change>2, p<0.05) and downregulation of 70 genes (blue dots, fold-change>2, p<0.05) in PHF3 ΔSPOC compared to WT. Drosophila S2 nuclei were used for spike-in normalization. (C,D) GO analysis of genes upregulated in PHF3 ΔSPOC and genes upregulated in both PHF3 KO and ΔSPOC shows enrichment of genes involved in neurogenesis. GSEA ‘Biological processes’ tool was used.

**Figure S7.**
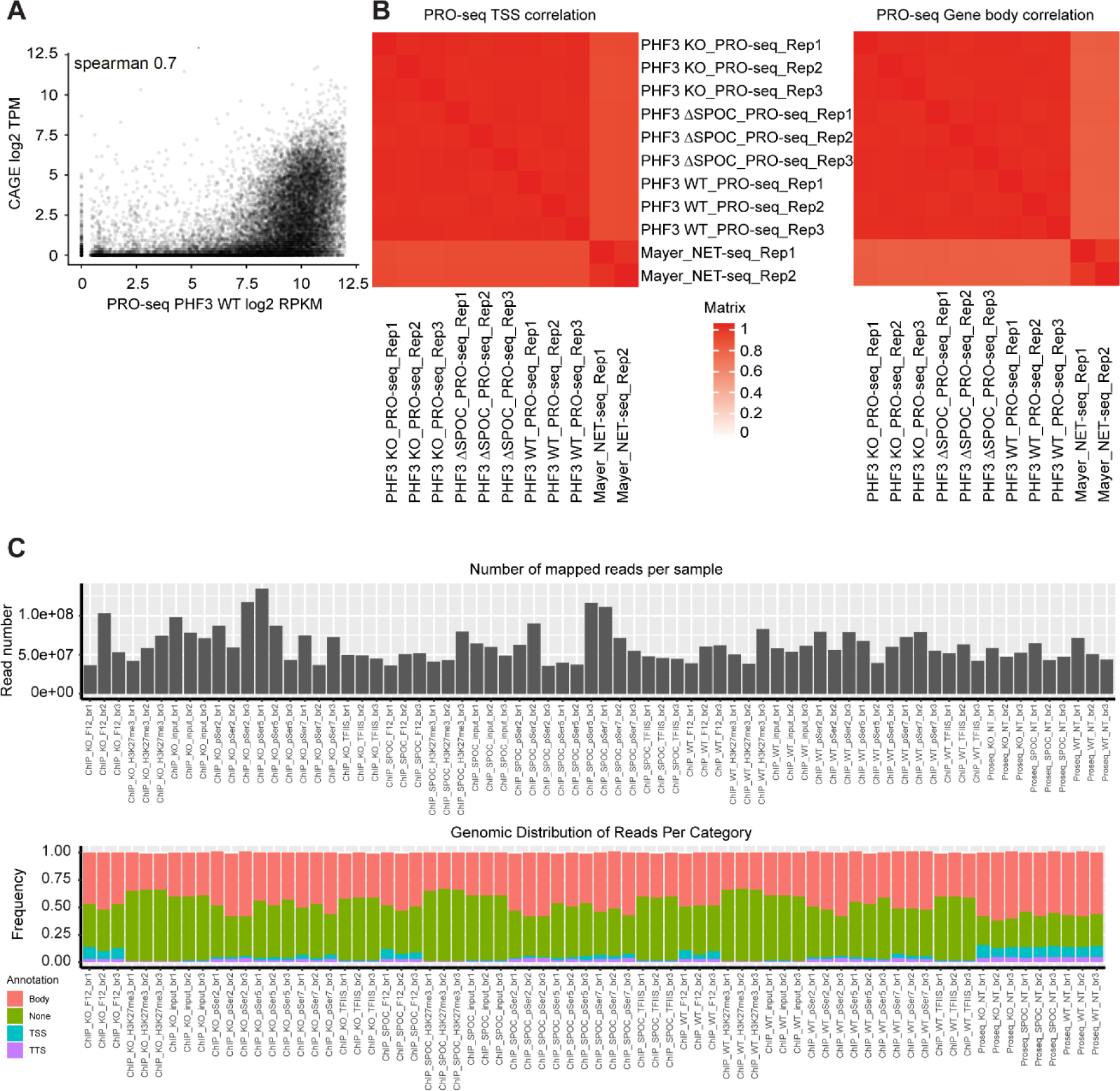
Validation of PRO-seq data analysis. (A) Comparison between our PRO-seq PHF3 WT and CAGE data. (B) Comparison between our PRO-seq data for PHF3 WT and KO, and published NET-seq data from HEK293 cells. (C) Distribution of ChIP-seq and PRO-seq data along the functional genomic features showing the absolute number of mapped reads (above) and the frequency of reads in different categories.

**Figure S8.**
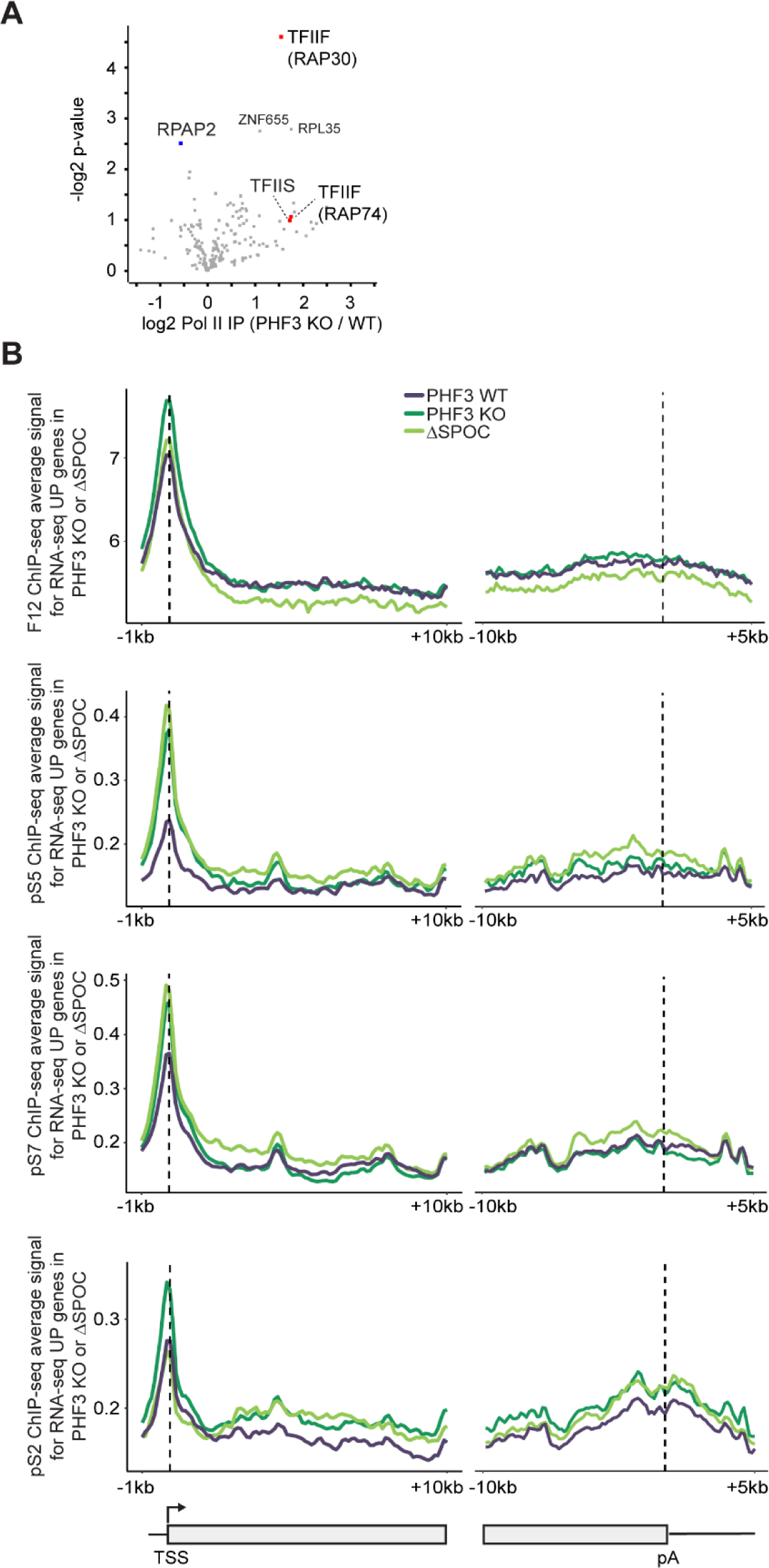
Pol II mass spectrometry and ChIP-seq analysis. (A) Volcano plot showing differential Pol II interactome in PHF3 KO/WT cells as judged by mass spectrometry analysis of Pol II F-12 pull-down samples (n=3). (B) Composite analysis of Pol II ChIP-seq distribution and signal strength in PHF3 WT, KO and ΔSPOC on TSS-gene body region and gene body-pA region for genes that were upregulated in RNA-seq.

**Figure S9.**
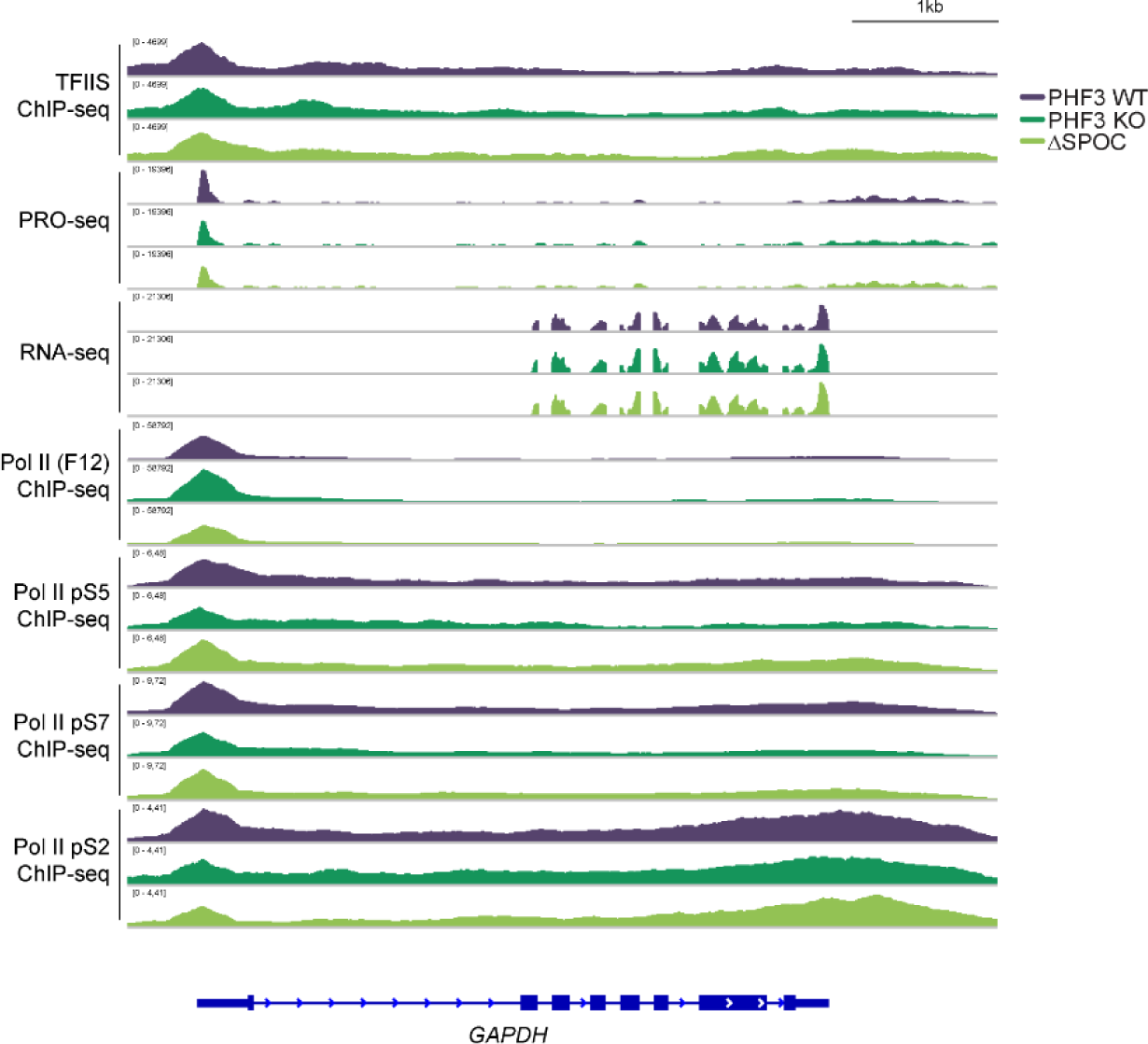
Genome browser snapshots showing TFIIS ChIP-seq, RNA-seq, PRO-seq, and Pol II ChIP-seq reads for GAPDH as a housekeeping gene

**Figure S10.**
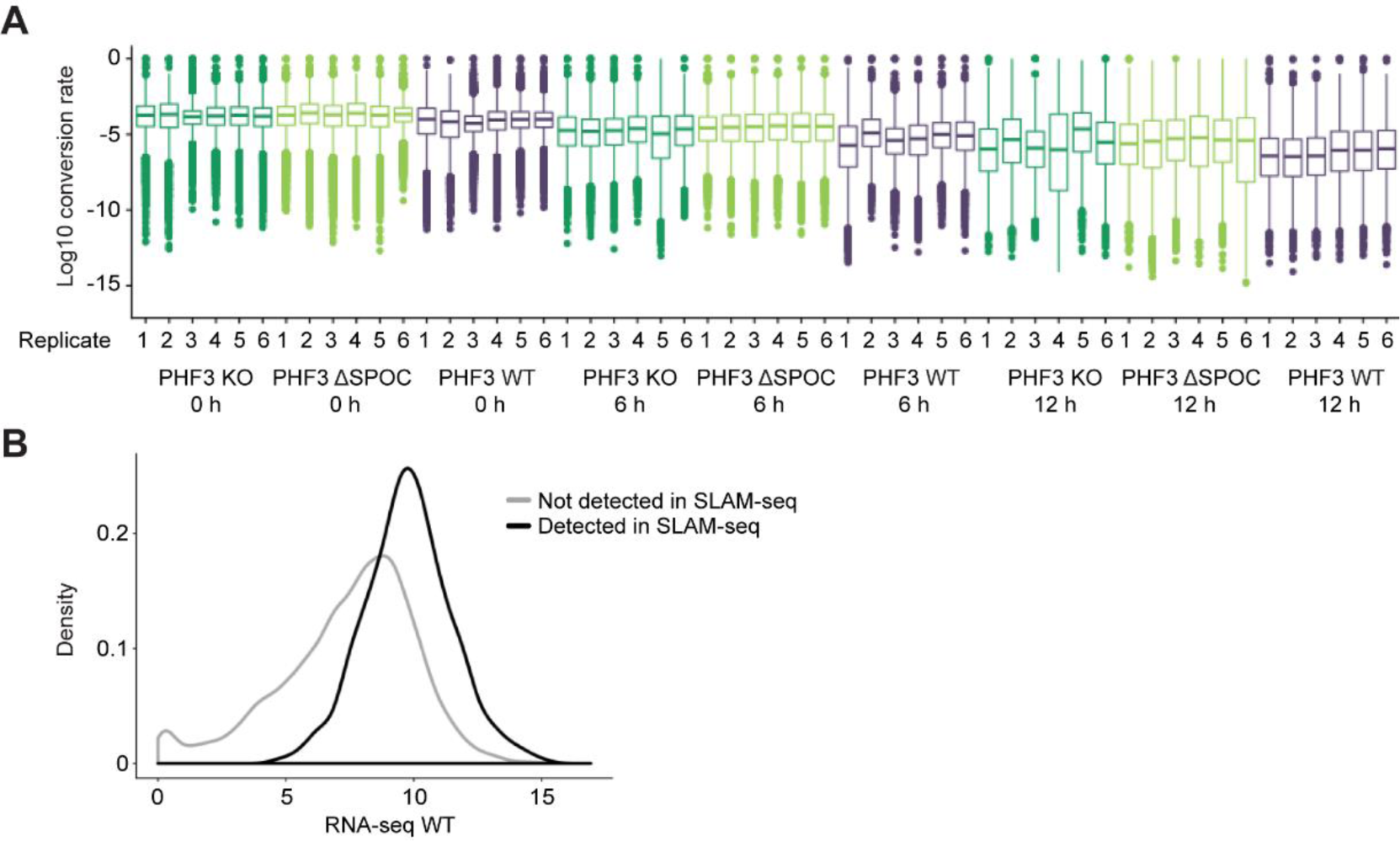
SLAM-seq analysis of mRNA stability in PHF3 WT, KO and ΔSPOC cells. (A) Conversion rate distributions per replicate for each genotype and condition. ‘0 h’ corresponds to 12 hours of s^4^U labelling, ‘6 h’ and ’12 h’ correspond to chase with uridine. (B) SLAM-seq captures mid/high expressed genes but not low-expressed genes.

**Figure S11.**
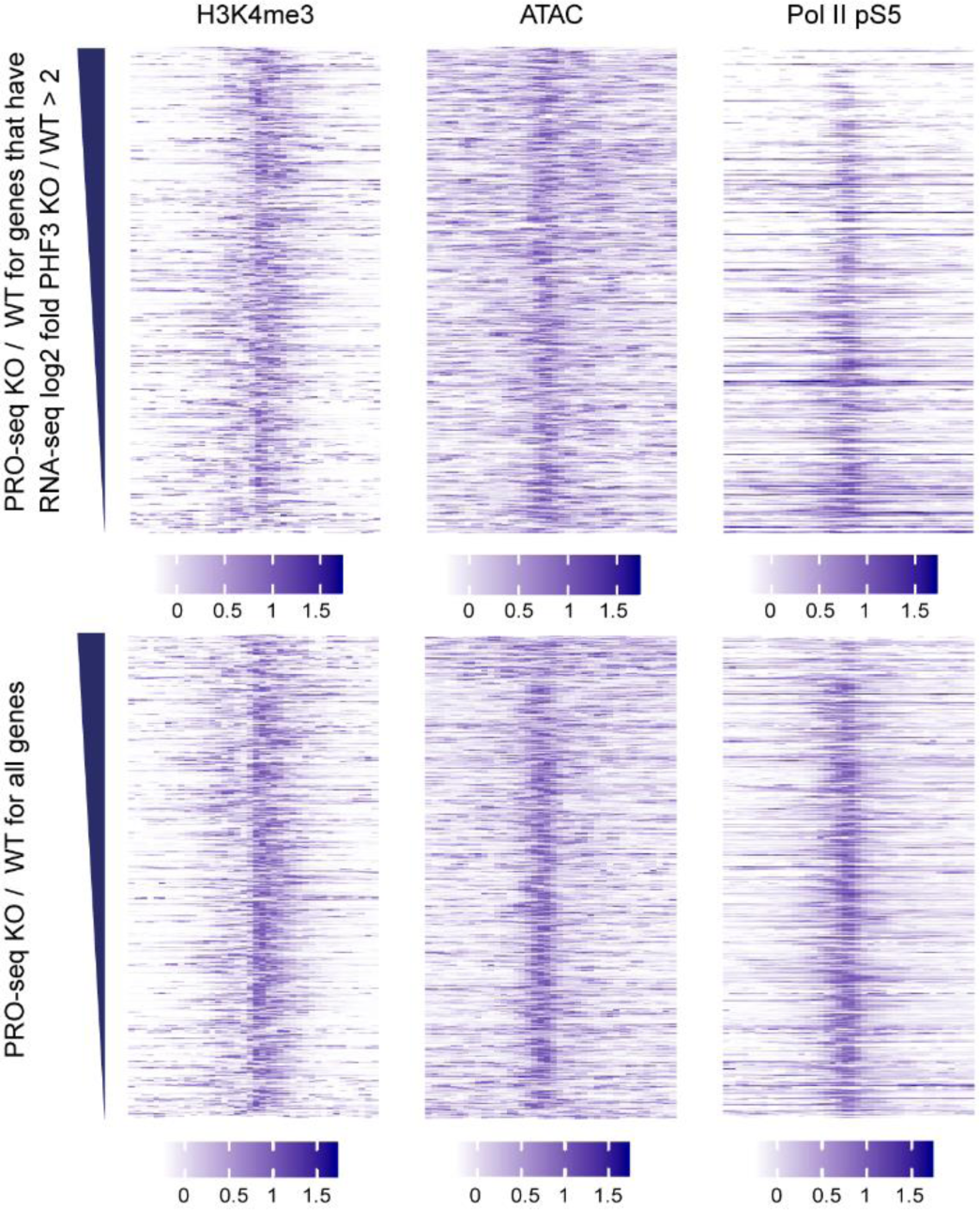
Promoter properties of genes that were upregulated in PHF3 KO and ΔSPOC. (A) Genes upregulated in PHF3 KO are enriched for H3K27me3 repressive mark but also exhibit features of open promoters such as H3K4me3, ATAC signal and Pol II pS5. The heatmaps show ChIP signal in the region of +/- 2kb around TSS. ATAC-seq data was from Array express ID: E-MTAB-6195; H3K4me3 data was from ENCODE ID: ENCSR000DTU. Prior to visualization, each signal in each TSS was scaled by dividing the corresponding values with the range (max value - min value). This was done for each experiment separately, in order to enable visualization of ChIP data on completely different scales.

**Figure S12.**
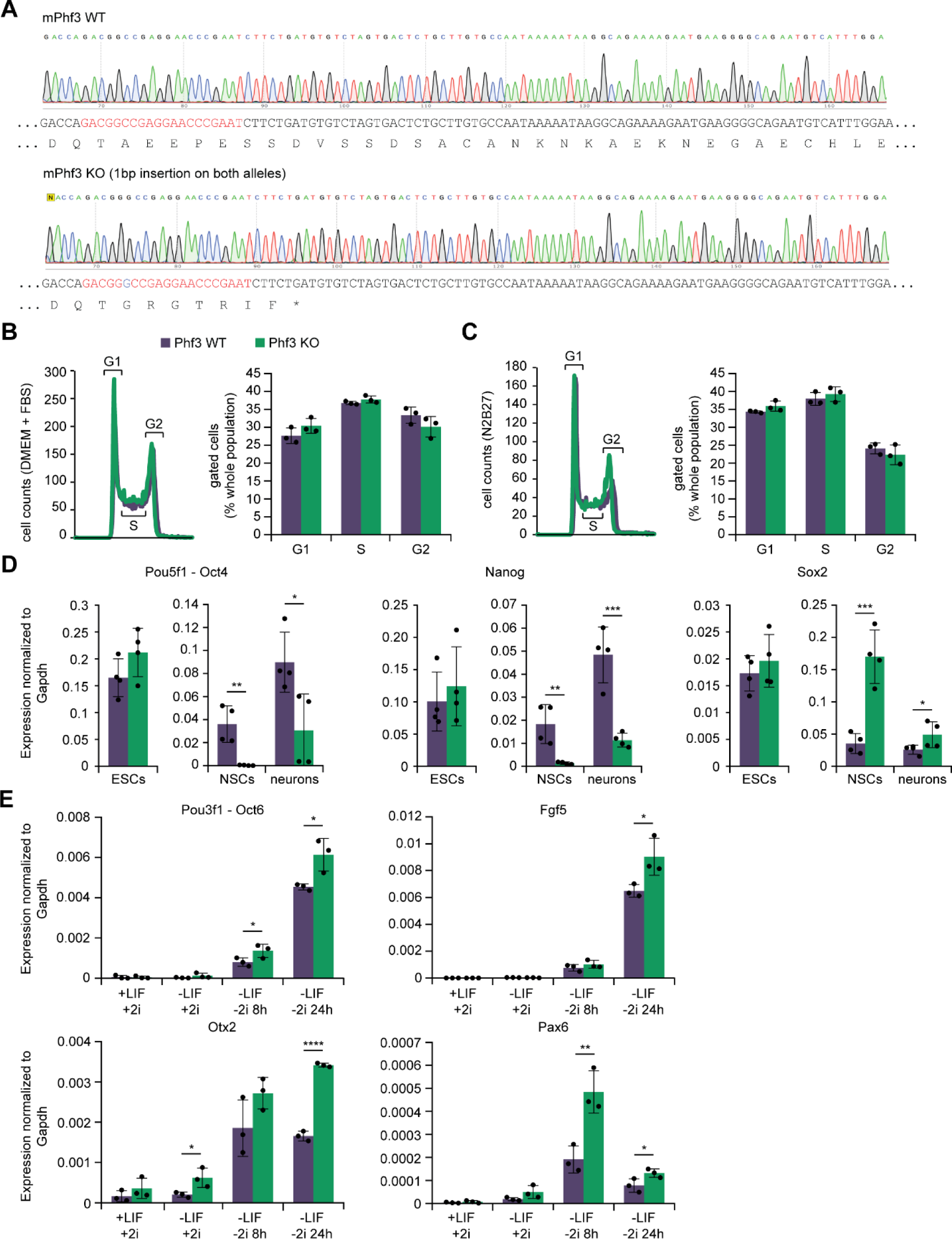
Generation of Phf3 KO mESCs and upregulation of early ectodermal markers in Phf3 KO ESCs. (A) Sequencing snapshots and sequence reconstruction of mouse Phf3 KO in ESCs. (B,C) FACS analysis of cell cycle distribution of propidium iodide-stained Phf3 WT and KO mESCs grown in (B) DMEM+FBS or (C) N2B27 supplemented with 2i/LIF. (D) RT-qPCR analysis of expression levels of stem cell markers Oct4, Nanog and Sox2 in Phf3 WT and KO during differentiation from mESCs to NSCs and neurons. (E) RT-qPCR analysis of changes in expression levels of early ectodermal markers Oct6, Fgf5, Otx2 and Pax6 during 24 h differentiation of Phf3 WT and KO mESCs.

**Figure S13.**
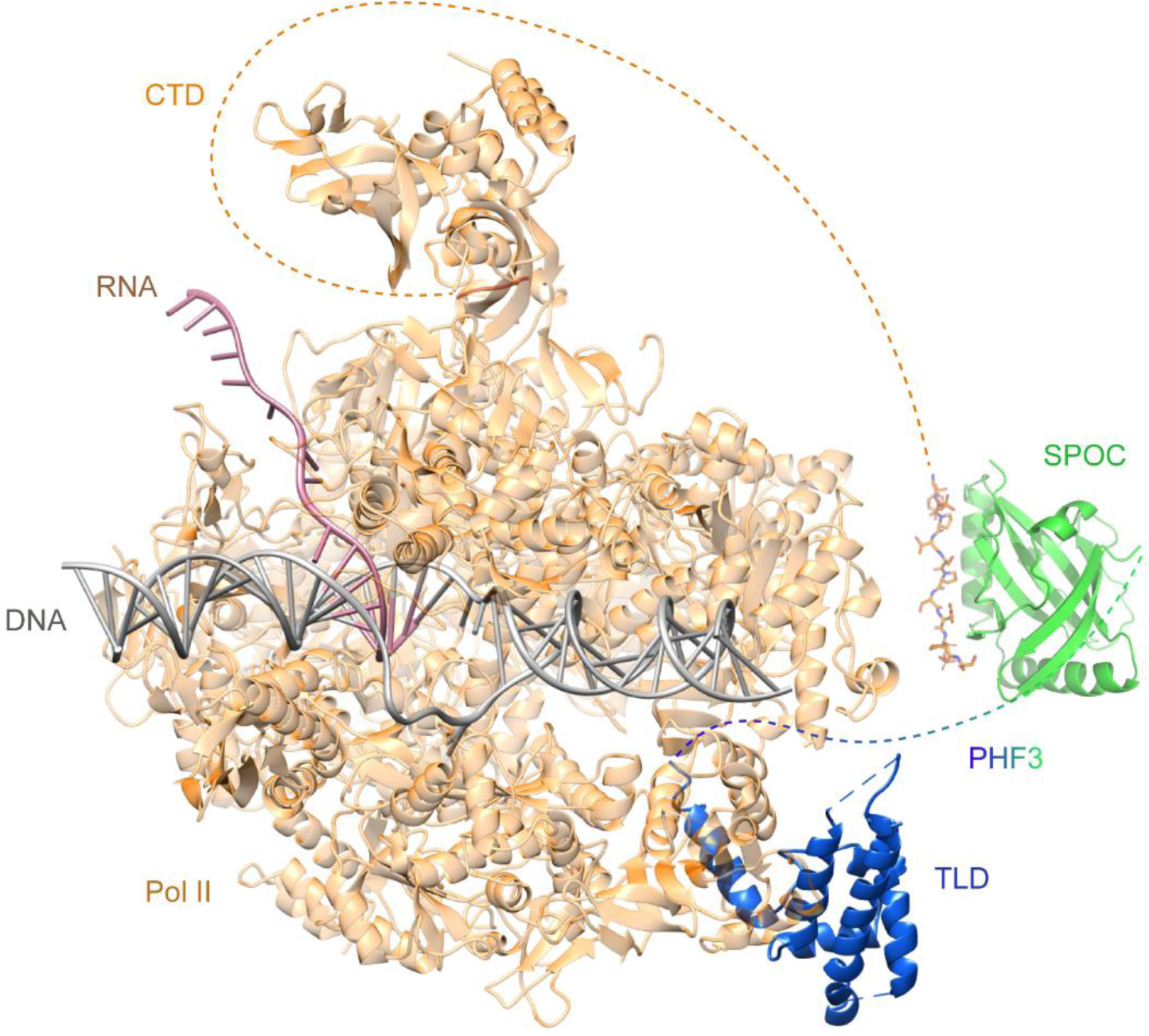
A structural model of PHF3 TLD-SPOC binding to Pol II. Yeast Bye1 TLD (4BY7) shown in blue was overlayed onto transcribing bovine Pol II (5OIK) shown in orange. PHF3 SPOC (green) in complex with 2xS2P CTD (orange) (6IC8) is connected with TLD and Pol II with dashed lines due to the absence of structural data for the disordered CTD and the linker between TLD and SPOC.

